# AlphaMut: a deep reinforcement learning model to suggest helix-disrupting mutations

**DOI:** 10.1101/2024.09.21.614241

**Authors:** Prathith Bhargav, Arnab Mukherjee

## Abstract

Helices are important secondary structural motifs within proteins and are pivotal in numerous physiological processes. While amino acids (AA) such as alanine and leucine are known to promote helix formation, proline and glycine disfavor it. Helical structure formation, however, also depends on its environment, and hence, prior prediction of a mutational effect on a helical structure is difficult. Here, we employ a reinforcement learning algorithm to develop a predictive model for helix-disrupting mutations. We start with a toy model consisting of helices with only 30 AA and train different models. Our results show that only a few mutations lead to a drastic disruption of the target helix. We further extend our approach to helices in proteins and validate the results using rigorous free energy calculations. Our strategy identifies amino acids crucial for maintaining structural integrity and predicts key mutations that could alter protein function. Through our work, we present a new use case for reinforcement learning in protein structure disruption.

## 2 Introduction

The structure of a protein is encoded in the sequence of amino acids. It is believed to be the free energetically the most stable state, or a state stuck at a local minimum with high barriers.[1]. The primary factors that govern the stability of a protein are the negative enthalpy change accompanying the stabilization of the structure due to hydrogen bonds and an increase in entropy resulting from the expulsion of water molecules from the peptide backbone[2]. Yet, the mechanism by which primary sequence information results in a thermodynamically stable structure remains largely unsolved[3, 4]. Recent advancements in deep learning techniques such as AlphaFold2[5], RosettaFold[6], and ESMFold[7], have greatly improved the accuracy of predicting protein structures. Whether these deep learning techniques “understand” the principles of protein folding is, however, debated.[4, 8, 9]

Helical moieties are the most common secondary structures in proteins, comprising 30% of all secondary structures in globular proteins. They play important physiological and biochemical roles, including signaling, recognition, and binding. The formation of hydrogen bonds (HBs) between specific backbone atoms leads to different types of helices, such as *α*, 3_10_, and *π* helices. *α*-helices, in particular, form HBs between the *i*-th and *i* + 4-th AA. Helices in proteins are favored due to a delicate balance between backbone hydrogen bonding and sidechain exposure to the solvent. While backbone hydrogen bonds are crucial for the initial nucleation of helices, side chains can increase or decrease stability and influence the dynamics of the helix through their solvent interactions, affecting both the nucleation and propagation phases of helix formation.[10]

Helix propensity (HP) refers to the likelihood or tendency of a specific AA to form an *α*-helical structure in proteins. It plays an important organizing role during the earliest stage of protein folding.[11]. Due to the Favorable interactions of alanine in the folded helical structure[12], alanine is considered to possess the highest HP, whereas glycine, which functions as a helix terminator, exhibits the lowest HP[13, 14]. The proline, which is excluded from this propensity calculation, is a helix breaker. Due to its cyclic side chain, it does not allow for favorable *ϕ* values for helix formation, causing kinks in the helical structure.[15]

However, HP is not uniform across the AA positions within a helix. Substitution of alanine with glycine destabilizes the helix more in the center than at the edge of a helix[16]. Likewise, the propensity for a polar AA is higher at the terminal region compared to the middle of the helix [17]. However, nonpolar amino acids show different behaviors; isoleucine and valine are more favorably found in the terminal region. In contrast, leucine and methionine are found more favorably in the center of the helix[18]. Helix propensity also differs between the cytosolic and membrane environments. *β*-sheet prompting amino acids such as isoleucine, valine, and threonine have been found in the transmembrane segments of alpha helices.[19] Similarly, proline, typically associated with a low helical propensity in cytosolic settings, exhibits a different behavior in membrane environments. [20] Aspartic acid mutations, meanwhile, can destabilize a helical peptide at the C-terminal end while stabilizing the helix at the N-terminal end[21].

Mutations and disruptions in helical structures are also crucial in disease contexts. Most disease-causing protein mutations are known to disrupt secondary structures[22]. Vihinen *et al*. showed that arginine is the most mutated amino acid in 44 disease-related proteins[22]. It has been shown that replacing polar amino acids can form non-native hydrogen bonds, leading to disruptions in protein structure and function and, thereby, disease conditions.[23] Disruptions in helix structures have also been linked to amyloid aggregation, causing diseases such as Parkinson’s and Cutzfeldt-Jakob disease (CJD) through misfolding of alpha-synuclein[24] and prion protein[25], respectively. Diseases such as non-Hogdkin lymphoma are caused by a conformational switch of the terminal *α*-helix to a *β*-strand caused by a D83V mutation[26]. Mutations in helices have also been shown to lead to fold switching. Mutations in the 11th amino acid of the Arc repressor of the bacteriophage P22 have been shown to be important in fold switching from two right-handed 3_10_-helices to wild-type *β*-ribbon [27]). Porter and coworkers demonstrated that amino acid mutations facilitated a switch from helix-turn-helix into a winged helix (helix-beta-sheet) in a family of 600,000 bacterial response regulator proteins using phylogenetic analysis along with ancestral sequence reconstruction [28]. From a purely biochemical point of view, understanding helix-disrupting mutations is particularly important since *α*-helices are structurally robust to mutations compared to other secondary structures[29].

While experimental investigations into helical propensity and disruption [30] involved synthesis and characterization, theoretical investigations include bioinformatic studies on structural datasets [22, 23] or molecular dynamics simulations post mutations[31]. However, both the experimental and molecular dynamics approaches are time-intensive. In addition, bioinformatic methods may lack generalizability if data are limited. Recent developments, including ESMFold and OmegaFold[32], which predict the structure of proteins from single sequences, have increased the speed of prediction by an order of magnitude compared to AlphaFold. Likewise, secondary structural contents can be easily computed using tools such as DSSP [33], P-SEA[34], and other machine learning techniques[35, 36]. A combination of these tools enable us to potentially perform large-scale mutational studies on proteins and study the effect on their structures, particularly helices, more efficiently.

Reinforcement learning (RL) is a subset of machine learning in which an agent learns to make decisions by interacting with an environment, receiving feedback, and refining its actions accordingly[37]. In the field of protein science, RL has been employed particularly in protein design tasks. EvoPlay, for instance, is a self-play reinforcement learning model that utilizes Monte Carlo tree search techniques to efficiently explore the design space, predicting peptide binders with increased affinity among other design tasks[38].

Here, we develop a generalized RL model **AlphaMut** that learns mutations detrimental to a helix in a given protein. Our approach integrates methodologies from protein structure prediction, secondary structure analysis, and reinforcement learning. Inspired by existing frameworks[38, 39], we conceptualize protein sequences as states, mutations as actions, and reward mutations that disrupt helices to some degree. Mutations that disrupt the helical structure are rewarded, while those that retain the helical structure are penalized. Through comprehensive training on helix datasets, we demonstrate the efficacy of our model in accurately predicting disruptive mutations within helices. We then proceed to derive a propensity index for the amino acids. The useful application of our approach comes from the extension of our model to helices within proteins, which allowed us to identify several sets of specific mutations in a protein that could potentially disrupt the helix or the entire structure. We validated our results through rigorous all-atom-free energy calculations. Thus, this work serves as a promising tool to identify essential AA in a protein whose mutations might disrupt the structural and functional integrity of the same.

## 3 Methods

### 3.1 Reinforcement Learning Framework

Reinforcement learning (RL) is a subset of machine learning that is developed to solve decision problems where the goal is for an agent to learn to perform actions that provide the best outcomes. The agent performs actions and receives feedback in the form of a reward from the environment, which helps to assess whether the action was beneficial, neutral, or detrimental. Through continuous interaction with the environment, the agent refines its policy, a function that maps states to actions, to improve its decision-making and performance, thereby optimizing the outcomes in a complex, often uncertain setting[37]. The rewards from the environment are often delayed, meaning that the benefits or consequences of certain actions might not be immediately apparent, requiring the agent to balance short-term rewards with longterm gains. Reinforcement learning has been applied successfully in games like Go[40] and Chess[41], where the outcome of the game is known only after several steps.

In our problem, the environment consists of protein sequences as states, point mutations as actions, and a measure of helix disruption as a reward. A sequence of *l* residues has ^*l*^*C*_*n*_ × 19^*n*^ possibilities for *n* mutations. The number of actions and outcomes in this problem is too large for one to exhaustively search, which is why we choose reinforcement learning to solve our problem. Reinforcement learning also handles delayed rewards effectively; a mutation that does not disrupt the helix immediately can, however, lead to disruption upon combination with the right set of future mutations.

There exist several reinforcement learning algorithms such as REINFORCE, PPO, DDPG, A2C, DQN and SAC.[42–46] One such algorithm, Proximal Policy Optimisation(PPO), which we use throughout our work, operates by improving the agent’s policy through small, stable updates. Through this strategy, PPO avoids drastic changes in the policy that could potentially destabilize learning, leading to poor performance. PPO accomplishes this by limiting the difference between old and new policies using a surrogate objective function that constrains the amount of policy updates.

In this work, we have studied two different environments for helix disruption: (i) helices in isolation (Helix-only model) and (ii) helices within the context of an entire protein (Helix-in-protein model). The first model serves as a model system to probe the helix-disrupting mutations and HP of amino acids, while the second model extends the approach to understanding the impact of the protein’s environment on helix disruption. A detailed description of both environments is given below.

#### States and Embeddings

To quantitatively represent protein sequences, we employ state-of-the-art embedding models such as ProtVec or ESM-2, which transform each sequence into a high-dimensional matrix, capturing the physicochemical properties and evolutionary information of amino acids. For the Helix-only mode, helix sequences of length 30 were represented as states using two different algorithms: (i) ProtVec[47], a continuous distributed NLP-based model, and (ii) ESM-2[7] (8 million parameters), a protein language model (Figure 1). For the Helix-in-protein model, helix sequences of length 15 were chosen and their corresponding protein sequence were represented as states using ESM-2[7] (8 million parameters). To obtain the states, we concatenated the ESM-2 representation of the helix and its corresponding protein (see Figure 3A).

**Figure 1.**
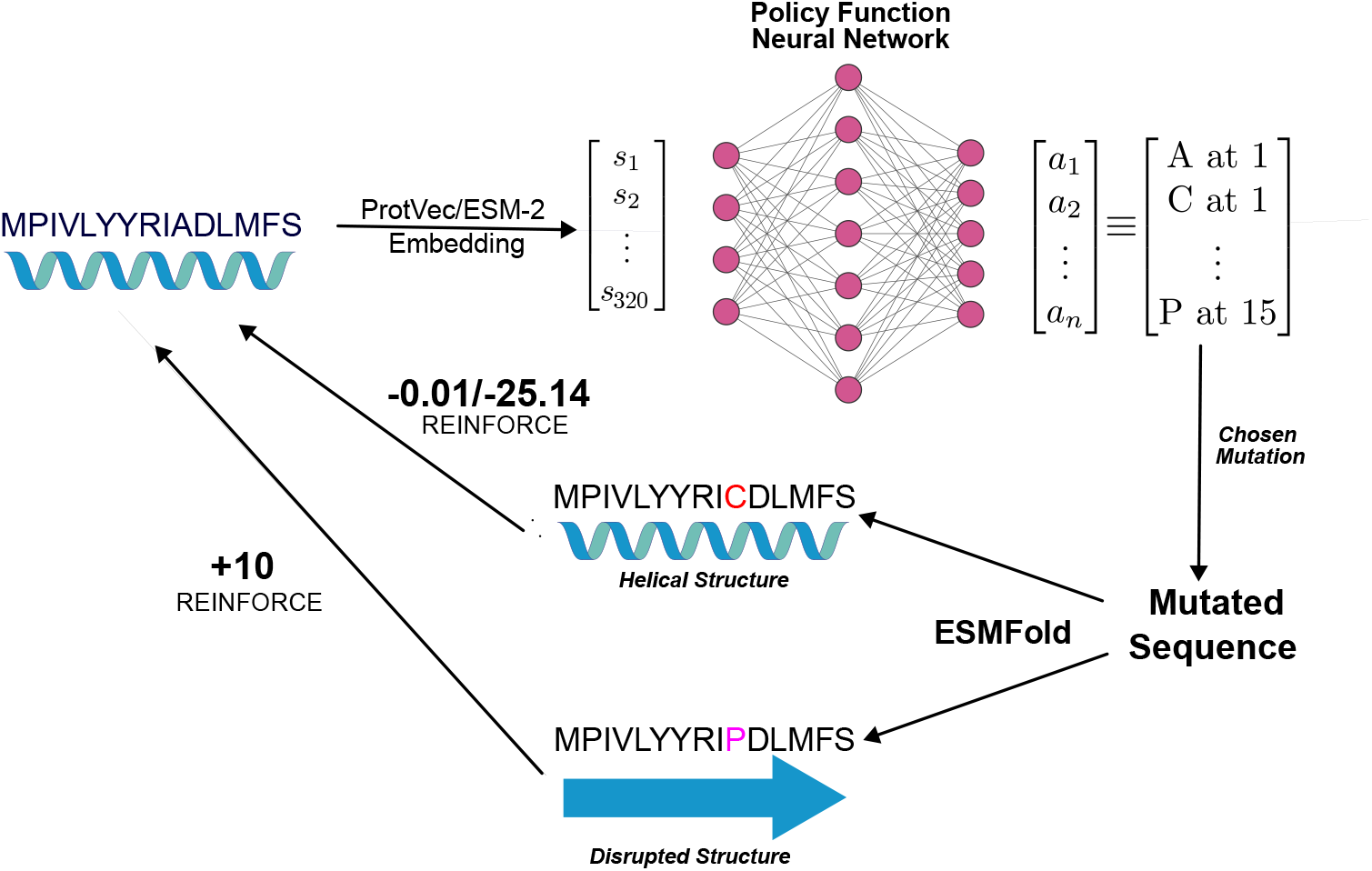
The Helix-only Reinforcement Learning Environment for a helix of 15 residues.

#### Actions

Actions were modeled as the set of all possible mutations. For both Helix-only and Helix-in-protein environments, we trained RL models with and without proline as an allowed mutation (since proline is an obvious helix-breaking amino acid). For the models trained with proline as a possible mutation, there were *l* × 20 actions, while for those trained without proline as a possible mutation, there were *l* × 19 actions, where *l* corresponds to the length of the helix – 30 for the Helix-only model, and 15 for the Helix-in-protein model.

#### Reward

The reward function in our environments was designed to encourage the reduction of helical content following a point mutation. Specifically, we defined successful disruption as a significantly high value of 70% reduction in helical content. To this end, we used ESMFold[7], a state-of-the-art method for structure prediction and the P-SEA algorithm[34] for secondary structure assignment to quantify the disruption of helix structures.

Upon selecting a mutation, percentage helical disruption (Δ) was calculated using the mathematical expression provided below:

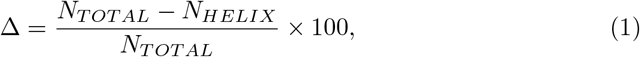

where *N*_*HELIX*_ is the number of residues part of a helix post mutation, while *N*_*TOTAL*_ is the total number of residues originally part of the helix. *N*_*HELIX*_ is calculated by summing all residues assigned as a helix by the P-SEA[34] algorithm. As described earlier, a successful disruption is defined when Δ reached greater than 70. For the Helix-in-protein model, Δ was calculated only for the helix part chosen to study the effect of mutations.

Since single-point mutations may not be sufficient to disrupt the helical structure, the RL model was allowed to execute up to *N*_*MAX*_ mutations within a single episode. A positive reward of +10 was provided if the helix was disrupted within *N*_*MAX*_ mutations. A penalty of −25 was imposed if the helix remained intact after *N*_*MAX*_ mutations. Additionally, a small negative reward was given whenever the helix was not disrupted upon taking an action (mutation), encouraging faster disruption. The mathematical description of the reward function, *R* is given below.

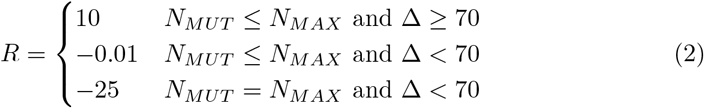

Here, *N*_*MUT*_ is the number of mutations. Episodes were terminated either when the helix was disrupted or if the number of mutations reached *N*_*MAX*_. This approach allows the model to experiment with different mutation patterns that can disruption to the helical structure. The RL environment, consisting of the states, actions and reward function, was constructed using the python package Gymnasium[48].

#### 3.1.1 Datasets of helices

RL models are trained with different initial states for each episode. In our environments, we choose a helix sequence as an initial state, and to do so, we created databases of helices for each environment.

##### Dataset for Helix-only model

The therapeutic peptides (TP)[49] database, a collection of all helical motifs in the Protein Data Bank (PDB), was used to build this database. The TP database was downloaded from https://dyn.life.nthu.edu.tw/design and filtered to gather helices with 30 amino acids. Duplicate sequences were dropped to obtain a non-redundant dataset of 2066 helices.

The dataset was further split into train-validation(90:10) sets to obtain 1874 helices for training and 192 for validation.

##### Dataset for Helix-in-protein model

Datasets were created by selecting helices of 15 amino acids in length from the TP dataset [49]. Subsequently, the entire RCSB-PDB sequence database [50] was downloaded to match and obtain complete protein sequences corresponding to the chosen helices. To avoid redundancy, we considered only a single entry among those entries where both the helix sequence and protein sequence were identical.

A 90:10 split of this dataset yielded 32,775 training entries and 3,643 validation entries. To facilitate easier training, a subset of these sequences was also created, containing proteins with 250 or fewer amino acids. This subset includes 8,794 training entries and 1,009 validation entries. All results presented correspond to this subset.

#### 3.1.2 Reinforcement learning algorithms

Proximal Policy Optimization (PPO)[45], Actor-Critic (A2C)[43], and Deep Q Network (DQN)[46] algorithms from the python package Stable Baselines3 [51] were selected to train the models. These algorithms were chosen for their suitability for environments with continuous state spaces and discrete action spaces.

Helix-only model was trained for 168,000 steps, while the Helix-in-protein model was trained for 190,000 steps. Two hyperparameters, *γ* and *β*, were varied to obtain the optimum model configurations. While *γ* controls importance given to later rewards, *β*, the entropy coefficient, regulates the diversity of actions predicted by the model.

#### 3.1.3 Validation method and metrics

Structures produced post mutations were saved, and the process was stopped when the value Δ exceeded the threshold for the model. The mutations suggested by the model for each protein in the validation set were then analyzed. To choose the best model, the following metrics were calculated.

##### Learning Efficiency (κ)

This metric indicates the average normalized reward, obtained by scaling the average reward using the following expression.

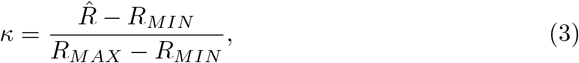

where *R*_*MAX*_ and *R*_*MIN*_ denote the maximum and minimum reward attainable per episode. This measure evaluates whether the model has learned effectively without overfitting the training dataset. A *κ* value close to 1 indicates better training, while a *κ* closer to 0 indicates overfitting. The 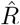 is the average reward obtained by running the model for 1000 episodes.

##### Mutation Entropy (S)

The frequencies of each amino acid predicted in episodes that successfully disrupted the structure were computed. The mutation entropy was calculated using the following equation.

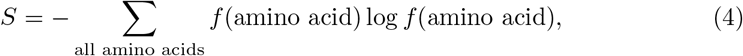

where *f* denotes the frequency and *S* reflects the diversity of the amino acid mutations proposed by the model. A higher *S* indicates more diverse amino acid mutations.

### 3.2 Free Energy Calculations

For validation, we calculated the free energetic stability of the wild-type and mutated helix in a protein environment. All-atom molecular dynamics (MD) simulations were performed with GROMACS version 2022.5[52]. Simulations were carried out in explicit water with the TIP-3P water model[53]. AMBER14SB[54] force field was used for the proteins. Every system was energy minimized using the steepest descent algorithm for 50000 steps. Subsequently, the NVT and NPT equilibration steps were performed for 5 ns and 10 ns, respectively. MD production run was performed for 10 ns. We utilized a velocity rescale [55] thermostat to control temperature and the Parrinello-Rahman barostat[56] to control pressure for NVT and NPT simulations.

Well-tempered metadynamics simulations were performed using GROMACS 2022.5[57] version patched with PLUMED 2.9.0[58]. We used ⟨Ψ⟩ as the collective variable (CV), defined as,

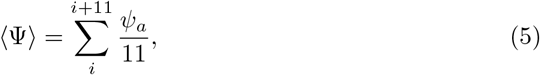

where *ψ* is the Ramachandran’s backbone torsional angle formed by *N*_*i*_−*Cα−C−N*_*i*+1_ atoms. *i* − 2 is the starting residue number of the helix. This allows us to exclude the terminal residues of the helix. For a helix, ⟨Ψ⟩ is around −40.5^*°*^, whereas it is between 120^*°*^ and 140^*°*^ for a *β*-sheet. We used this CV to study helix to non-helix transitions, as it was shown to be a good collective variable for the *α*-helix to *β*-sheet transition[59].

## Results

AlphaMut is a a set of reinforcement learning models that suggest mutations that can disrupt helices. While the Helix-only models disrupt an isolated helix, the Helix-in-protein models disrupt helices in a protein. We describe results from both models.

### 3.3 Helix-only Model

We first constructed an RL environment to disrupt a helix in isolation, i.e., independent of the protein of which it could be a part (Figure 1). To this end, we trained our model on data sets of helices with 30 amino acids. After experimenting with multiple algorithms (Advantage Actor-Critic (A2C), Proximal Policy Optimisation(PPO), and Deep Q-Network(DQN)), we chose PPO to train our model. DQN failed to converge(Figure S1), and while A2C converged quickly, it showed poor validation metrics (Table S1). PPO showed the best training among all algorithms, reflected in its convergence (Figure S1) and validation metrics(Table S2).

With PPO as our RL algorithm, we trained our model using two embedding types: from ProtVec[47], an unsupervised data-driven distributed representation, and ESM-2[7], a protein language model. We also varied the hyperparameters *γ* (which penalizes later rewards) and the *β* (which encourages diverse actions). To evaluate these approaches, we used our validation split to calculate two validation metrics: (i) mutation entropy (*S*)(a measure of the amino acid diversity of predicted mutations) and (ii) learning efficiency *κ* (a measure of model learning effectiveness). High values in both metrics indicate a well-trained model that predicts a variety of amino acid mutations. The validation metrics for all strategies are given in Table S2. Upon convergence of the PPO-trained models (Figure S1), we evaluated their corresponding validation metrics and selected the model with the parameters highlighted in Figure S3. Visual inspection of the disrupted structures revealed kinks in the helix, characteristic of proline mutations (Figure S3). To quantify this observation, we analyzed the frequency of mutations as shown in Figure 2A. The model predicted proline mutations 53% of the time, followed by glycine. This result is expected because both amino acids are known helix breakers, indicating that our model accurately represents a physical phenomenon rather than generating random predictions. Simultaneously, our method also provides data on the tendency of different amino acids to remain in a helical structure (less prone to mutation) and can thus be equated to helix propensity. Therefore, with information about predicted mutations against present amino acids, we propose a new approach to calculate the helical propensity of amino acids by dividing the frequency of occurrence of an AA in the database by the frequency of mutation of that AA(see Figure 2B). This approach is different from methods in the literature, which primarily consider the frequency of occurrence of an AA in existing helical datasets[49, 60], while it is similar to another method [61] that considers mutations while calculating propensities, albeit for amino acid groups. Our analysis shows that leucine has the highest propensity, while proline has the lowest. Correspondingly, we also obtain a matrix of which amino acids currently in the helix are important for the helix. (Figure S4)

**Figure 2.**
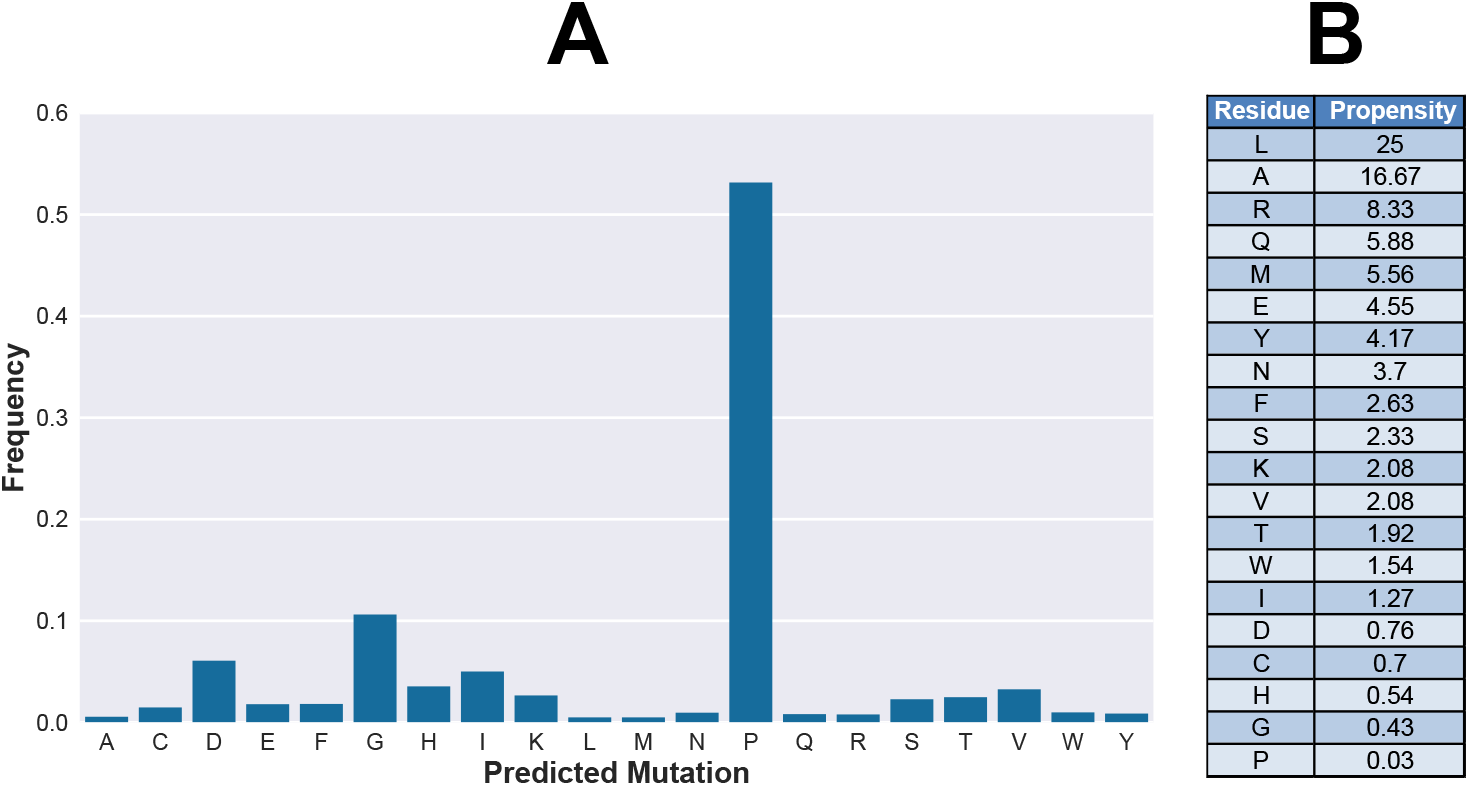
(A) Frequencies of the different amino acids predicted as mutations by the chosen Helix-only model. (B) Helix propensity derived from the validation studies.

While we generate different models with varying hyperparameters, we observe that the model predicts proline very frequently (53%), a term referred to as mode collapse in RL. To circumvent this, we removed proline as one of the options(mutations) in the environment and trained the model using the PPO algorithm. We observe that the trained model was successful in breaking helices without proline. However, it showed a lower learning efficiency. Although the proline-trained model was successful *>* 95 % of the time in disrupting the helix, the model without proline was successful at predicting disrupting mutations only around 80% of the time (see Table S2). Nevertheless, we analyzed the mutations predicted by the model and show that, in this case, the model mode collapses to glycine (see Figure S2).

### 3.4 Helix-in-protein Model

The largest set of experimentally characterized helices we can obtain is from the RCSB-PDB[50] through the therapeutic peptides database[49]. However, while these helices can be stabilized inside the protein by tertiary interactions, it is not known if they can exist as helices independently, i.e., without their protein environment. To investigate the influence of the protein environment on helix disruption, we constructed an RL environment to disrupt the helical structures in proteins (Figure 3A). Unlike the previous environment, we provide information on both the protein sequence and the helix sequence to provide context of the position of the helix within the protein environment. Like earlier, after the entire protein was modeled, we calculated the reward by considering the helical content of only the helix.

**Figure 3.**
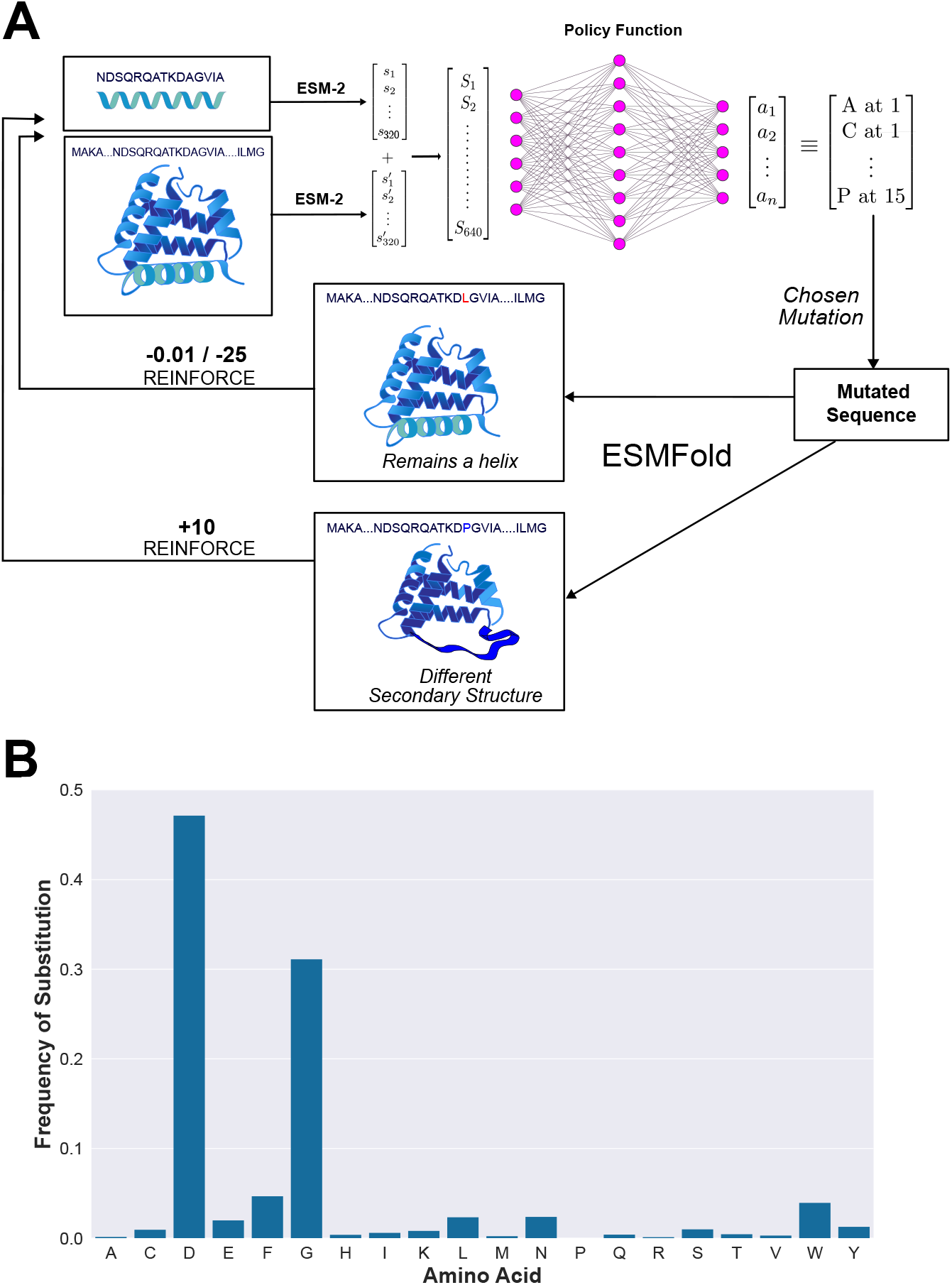
(A) Schematic of the RL Environment for the Helix-in-protein environment and (B) Barplot of the frequency of predicted substitutions.

Upon training this model with *M*_*MAX*_ = 5, 7, we plotted the frequencies of each predicted mutation and observed that this model also mode-collapsed to proline, and the mode collapse was more severe. While in the Helix-only model, proline was predicted as the mutation about 60 % of the time, here, proline is predicted as the mutation *>* 95 % of the time(see Figure S7). The results of these validations are given in Table S3. We also calculated the helical propensity obtained from this model and compared it with the values obtained from the Helix-only model. We observe that the helical propensity scales of amino acids with and without the protein environment are varied with a Spearmann correlation coefficient (*ρ*) of 0.43. It is thus important to always consider the environment in which the helix is present when trying to mutate it. Propensities in isolated helices and helices in proteins do not correlate well with each other. Therefore, an AA mutation that is relevant in an isolated helix may lose its importance when that helix is part of a protein, contrary to what is generally accepted in the literature[62].

To mitigate the mode collapse effects of proline, we trained our model without proline as an available mutation. We trained our model on *M*_*MAX*_ = 13 for 168000 steps. The convergence plot of this is shown in Figure S8. Our model yielded a learning efficiency of 0.54 indicating that it predicts a disrupting set of mutations 1 in every 2 times. We find a mutation entropy of 1.57, indicating that the diversity of mutations is higher compared to the model with proline. Particularly, we observe that aspartic acid is often predicted (47%) as a disrupting mutation. Likewise, glycine, a known helix-disrupting mutation, is also predicted (see Figure 3B)

To understand the mutations and the corresponding structures associated with them, we inspected the ESMFold-generated structures of different important proteins. Presented here in Figure 4 are mutated helix structures of frataxin and protein-L. Frataxin is a protein whose mutations are implicated in Frederic’s ataxia[63, 64] and protein-L is an immunoglobulin binding protein studied particularly in the context of protein design[65]. The effects of mutations on the structure of both proteins have been extensively studied through MD simulation, and therefore these proteins serve as ideal systems to test our model[66–68]. From the predictions of our model, frataxin presents a lot of triple mutants whose helices are disrupted. In protein-L, it is interesting to note the helix-to-sheet transition happening across mutations. The predicted mutant structures of other proteins such as insulin, hemoglobin, myoglobin, zinc finger protein, and HSP-90 are provided in Section 8 of the SI as some representative systems of interest.

**Figure 4.**
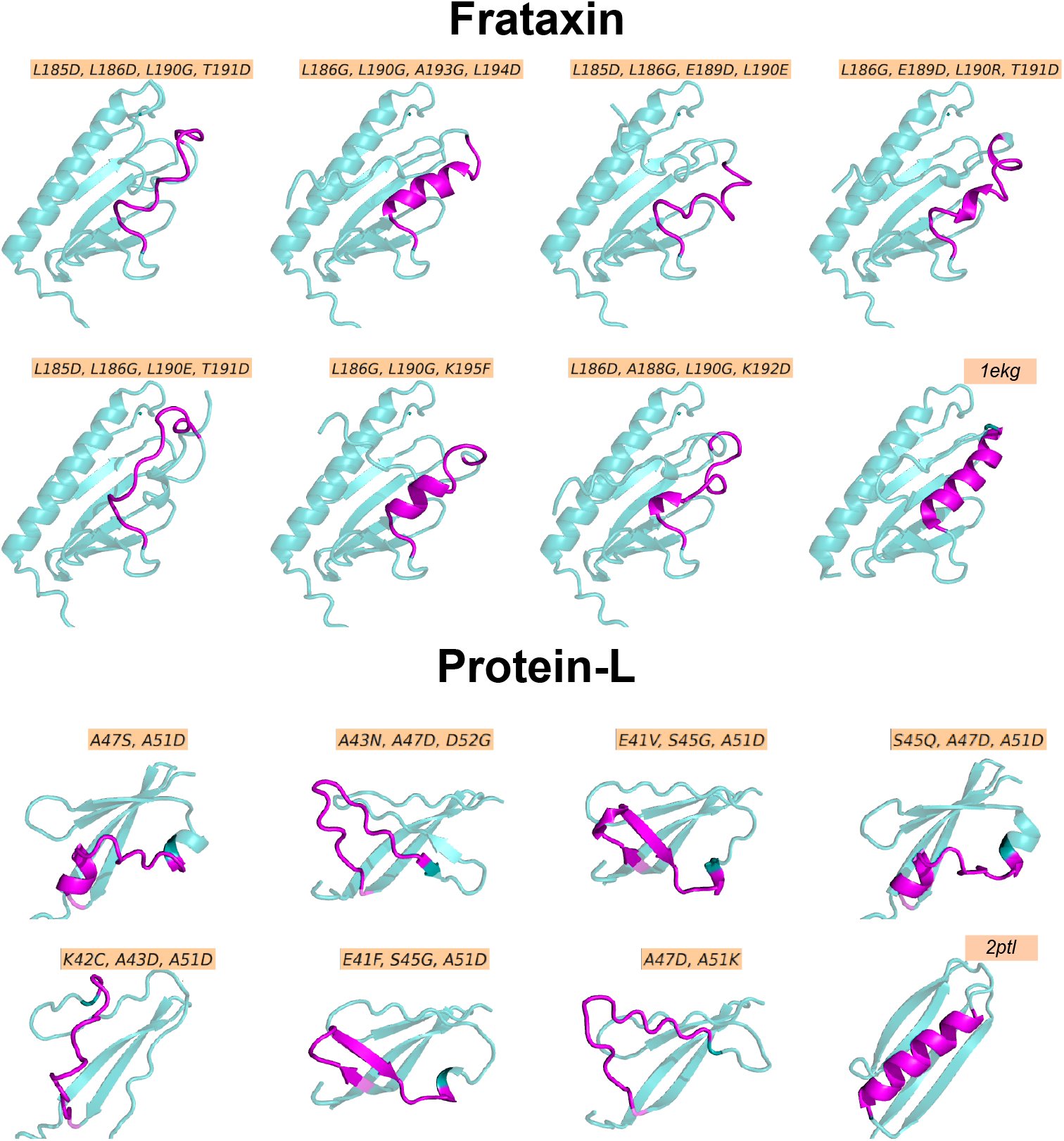
Structures of mutated proteins, Frataxin and Protein-L, along with their respective wild type structures and PDB IDs (right corner). Mutations introduced are indicated in an orange box above the respective structure.

To further validate if the mutations predicted by our model indeed resulted in disruption, we performed well-tempered (WT) metadynamics simulations to study the stability of the helical region in the wild-type and mutant. WT metadynamics is a computational technique that can be used to assess the stability of protein structures by exploring their free energy landscapes against a particular collective variable (CV). Since we are interested in helix disruption, we defined a CV, average-psi (⟨Ψ⟩), which captures the average value of the dihedral angle *ψ* of the helix we intend to disrupt. By enhancing the sampling along this CV, we captured various metastable states of both wild-type and mutant proteins. The success of our model depends on mutation prediction that lead to a deviation of the free energetically most stable state of the helical region from the ⟨Ψ⟩ value of ≈ − 40.5°, typical for a helix. To avoid any biases imposed by ESMFold, we modeled the point mutations using MODELLER[69] and performed the WT metadynamics simulations of for wild-type and mutant of frataxin and protien-L until convergence (see Figure S14,Figure S15). The free energy surfaces (FES) of all systems along the CV ⟨Ψ⟩ are provided in Figure 5. For the wild-type frataxin protein, we observe the most stable state around -40 °(Figure 5A). This value is very close (≈ − 40.5°) to ⟨Ψ⟩ for an ideal *α* helix, also confirmed by visual inspection of the structures. Based on the predictions of the model on the helix 182-196 of frataxin, we used a tetramutant L186D, A188G, L190G, and K192D. The most stable meta-stable state for this mutant is at ⟨Ψ⟩ ≈ +47 °, which corresponds to a random coil structure (Figure 5A). Aspartic acid mutations leading to disruption have rarely been studied in the literature, and helix disruption due to an aspartic acid mutation has been attributed to its interaction with the helix dipole[21]. The wild-type protein-L exhibits two stable metastable states: one around -40°, corresponding to a helical structure, and another around +10°, which does not form a helix. While these two states exhibit approximately the same stability, they are separated by a high free energy barrier of 40 kJ/mol. The protein-L mutant simulations are for the model-predicted triple mutant — E41F, S45G, A51D for residues 40-54. We observe the most stable state around ⟨Ψ⟩ ≈ 17°, which is a partial coil-helix structure toward the C-terminal end (Figure 5B). While other meta-stable minima are seen, they are not as stable as the minima at 17°. Simulations of both frataxin and protein-L highlight the critical role of aspartic acid mutations in structural disruption. Analysis of the free energy surface (FES) shows that the model-predicted mutations lead to a shift in the most stable metastable state from a helical to a non-helical conformation, validating the predictions made by AlphaMut.

**Figure 5.**
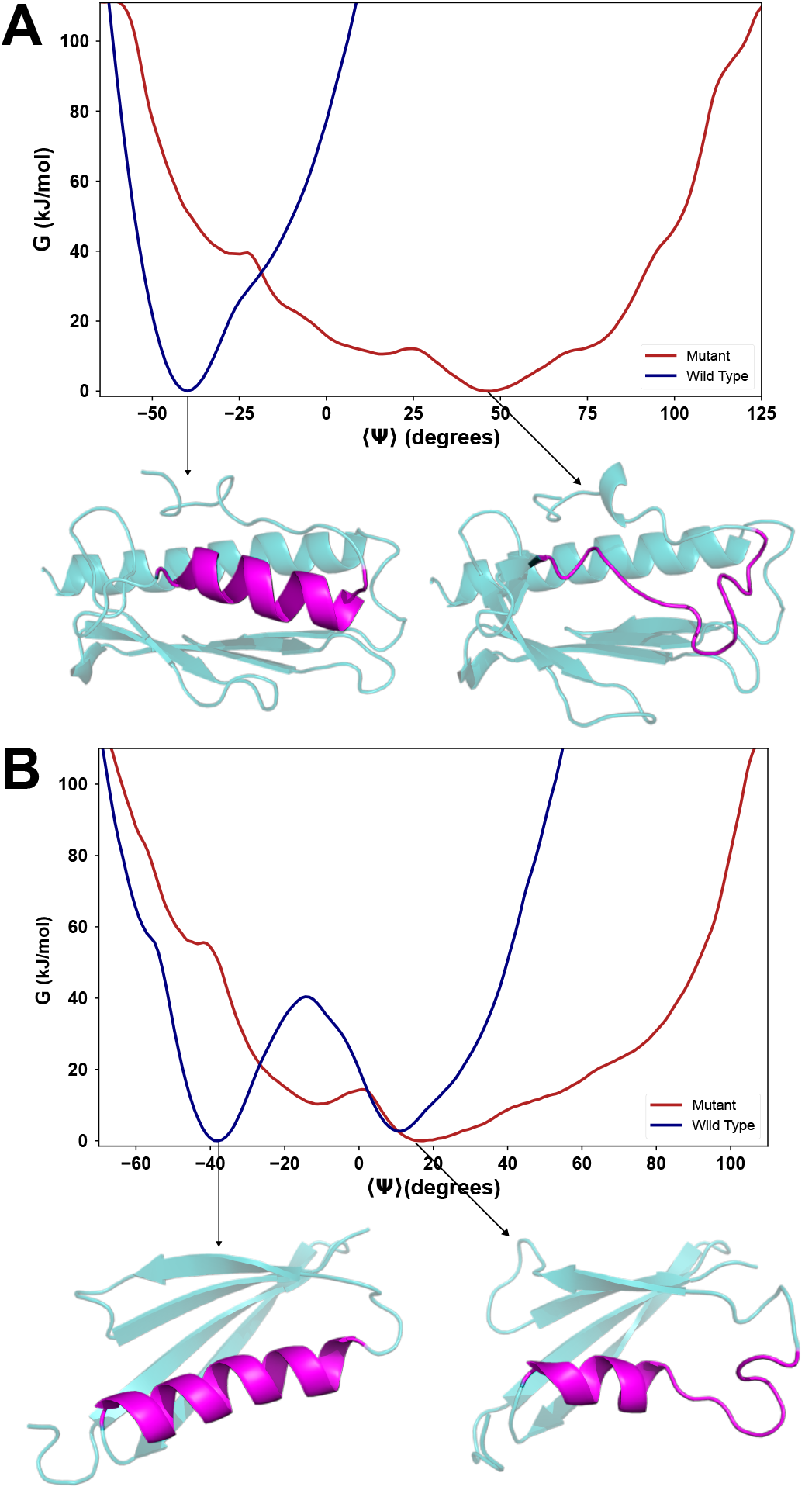
Free Energy Diagrams of wild-type and mutant Structures in (A) frataxin and (B) protein-L, with their minima structures indicated by an arrow. Mutations in the respective proteins are indicated in an orange box. The residues chosen for disruption are highlighted in magenta.

## 4 Conclusion

Helices are important secondary structures found abundantly in proteins. The disruption of helices has been widely studied through experiments and simulations. However, there is no general model for helix disruption. In this study, we developed a reinforcement learning model, AlphaMut, to predict mutations that would disrupt a helix either in isolation or within a protein environment. Expanding on previous literature, we calculated the helix propensity, not from the AA present in a helix as is usually done, but from the mutations that would cause disruption of the helix. We find that leucine has the highest helix propensity and proline has the lowest. This gives us an idea about which amino acids to target to break a helix, given a helix-breaking mutation. We also demonstrated that helical disruption is possible without using proline, but the average number of mutations required for disruption increases. After applying our approach to independent helices, we extend it to disrupting helices within a protein, to account for also the effect of tertiary interactions that stabilize a helix. This model almost always proposed proline as the mutation to disrupt helices.

We further trained our model without proline as a possible mutation and observed that aspartic acid and glycine break the helix the most frequently. While the helix-breaking properties of glycine are well-known because of its high conformational flexibility, there is hardly any literature citing aspartic acid as a helix-breaker. More mechanistic studies are needed to understand the role that aspartic acid plays in helix disruption. However, our results should not be viewed only in the context of the helix propensity of a single amino acid. Our model takes into account the entire sequence to predict the relevant set of mutations (not just one) that lead to the breaking of the helix. We further applied our model to a few important proteins that suggested a set of helix-breaking mutations. To validate the veracity of such mutations, we calculated the free energy surface of the structural stability of the helix for two proteins, frataxin and protein-L, using explicit water all-atom, free energy calculations using well-tempered metadynamics simulations. The results show that the most stable free energy for the wild type corresponds to helix, while for the mutant the stability shifts away from the helix, as expected.

The prediction of mutation effects on proteins has been extensively explored in the literature. Models such as AlphaMissense[70] and Missense3D[71] predict the pathogenicity of point mutations while other programs such as ThermoMPNN[72], RaSP[73], and DeepDDG[74] estimate stability changes upon mutations. Our procedure, on the other hand, accomplishes the same with an opposite approach, i.e., by predicting mutations that disrupt the helix. In addition, the results of our study demonstrate the effectiveness of the RL algorithm in manipulating protein structures.

AlphaMut is useful in assessing the stability of helical regions within proteins. Additionally, it would help identify appropriate mutations that specifically target helical structures, providing a deeper understanding of the protein’s structure and function. Most importantly, our approach and the model offer a systematic approach that goes beyond intuitive guesses or random mutation trials in the study of sequence-structure-function relationships in proteins through mutation-based assays.

## 5 Associated Content

The AlphaMut reinforcement learning environment and model have been provided on a GitHub repository - https://github.com/prathithbhargav/AlphaMut. A Colab notebook that runs the model is also available in the same repository.

## 6 Acknowledgement

The authors acknowledge the support and the resources provided by PARAM Brahma Facility under the National Supercomputing Mission, Government of India at the Indian Institute of Science Education Pune. The authors acknowledge the computational facility provided by the data science cluster of IISER Pune. PB acknowledges the KVPY fellowship from the DST, Government of India. PB acknowledges Ashish Kumar and Sonali Jadhav, IISER Pune for discussions and help in running metadynamics simulations.

## References

1. Solomatin, S. V., Greenfeld, M., Chu, S. & Herschlag, D. Multiple native states reveal persistent ruggedness of an rna folding landscape. Nature 463, 681–684 (7281 2010).

2. Dragan, A., Crane-Robinson, C. & Privalov, P. L. Thermodynamic basis of the α-helix and DNA duplex. European Biophysics Journal 50, 787–792 (2021).

3. Moore, P. B., Hendrickson, W. A., Henderson, R. & Brunger, A. T. The protein-folding problem: Not yet solved. Science 375, 507–507. eprint: 10.1126/science.abn9422. https://www.science.org/doi/abs/10.1126/science.abn9422 (2022).

4. Chen, S.-J. et al. Protein folds vs. protein folding: Differing questions, different challenges. Proceedings of the National Academy of Sciences 120, e2214423119 (2023).

5. Jumper, J. et al. Highly accurate protein structure prediction with AlphaFold. en. Nature 596. Number: 7873 Publisher: Nature Publishing Group, 583–589. issn: 1476-4687. https://www.nature.com/articles/s41586-021-03819-2) (2023) (Aug. 2021).

6. Baek, M. et al. Accurate prediction of protein structures and interactions using a three-track neural network. Science 373, 871–876 (2021).

7. Lin, Z. et al. Evolutionary-scale prediction of atomic-level protein structure with a language model. Science 379, 1123–1130 (2023).

8. Chang, L. & Perez, A. AlphaFold2 knows some protein folding principles en. Aug. 2024. 10.1101/2024.08.25.609581v1 (2024).

9. Outeiral, C., Nissley, D. A. & Deane, C. M. Current structure predictors are not learning the physics of protein folding. Bioinformatics 38, 1881–1887 (2022).

10. Miller, S. E., Watkins, A. M., Kallenbach, N. R. & Arora, P. S. Effects of side chains in helix nucleation differ from helix propagation. Proceedings of the National Academy of Sciences 111, 6636–6641. eprint: 10.1073/pnas.1322833111. https://www.pnas.org/doi/abs/10.1073/pnas.1322833111 (2014).

11. Aurora, R., Creamer, T. P., Srinivasan, R. & Rose, G. D. Local interactions in protein folding: lessons from the α-helix. Journal of Biological Chemistry 272, 1413–1416 (1997).

12. Scott, K. A., Alonso, D. O. V., Sato, S., Fersht, A. R. & Daggett, V. Conformational entropy of alanine versus glycine in protein denatured states. en. Proceedings of the National Academy of Sciences 104, 2661–2666. issn: 0027-8424, 1091-6490. 10.1073/pnas.0611182104 (2024)(Feb. 2007).

13. Nick Pace, C. & Martin Scholtz, J. A Helix Propensity Scale Based on Experimental Studies of Peptides and Proteins. Biophysical Journal 75, 422–427. issn: 0006-3495. https://www.sciencedirect.com/science/article/pii/S0006349598775290 (2024) (July 1998).

14. O’Neil, K. T. & DeGrado, W. F. A thermodynamic scale for the helix-forming tendencies of the commonly occurring amino acids. Science 250, 646–651 (1990).

15. Piela, L., Némethy, G. & Scheraga, H. A. Proline-induced constraints in α-helices. Biopolymers: Original Research on Biomolecules 26, 1587–1600 (1987).

16. Chakrabartty, A., Schellman, J. A. & Baldwin, R. L. Large differences in the helix propensities of alanine and glycine. Nature 351, 586–588 (1991).

17. Petukhov, M., Uegaki, K., Yumoto, N. & Serrano, L. Amino acid intrinsic -helical propensities III: Positional dependence at several positions of C terminus. Protein Science : A Publication of the Protein Society 11, 766–777. issn: 0961-8368. https://www.ncbi.nlm.nih.gov/pmc/articles/PMC2373540/ (2024) (Apr.2002).

18. Petukhov, M., Muñoz, V., Yumoto, N., Yoshikawa, S. & Serrano, L. Position dependence of non-polar amino acid intrinsic helical propensities. Journal of molecular biology 278, 279–289 (1998).

19. Li, S.-C. & Deber, C. M. A measure of helical propensity for amino acids in membrane environments. Nature structural biology 1, 368–373 (1994).

20. Li, S.-C., Goto, N. K., Williams, K. A. & Deber, C. M. Alpha-helical, but not beta-sheet, propensity of proline is determined by peptide environment. Proceedings of the National Academy of Sciences 93, 6676–6681 (1996).

21. Huyghues-Despointes, B. M., Scholtz, J. M. & Baldwin, R. L. Effect of a single aspartate on helix stability at different positions in a neutral alanine-based peptide. Protein Science : A Publication of the Protein Society 2, 1604–1611. issn:0961-8368. https://www.ncbi.nlm.nih.gov/pmc/articles/PMC2142265/ (2024)(Oct. 1993).

22. Khan, S. & Vihinen, M. Spectrum of disease-causing mutations in protein secondary structures. en. BMC Structural Biology 7. Publisher: BMC, 56. https://www.ncbi.nlm.nih.gov/pmc/articles/PMC1995201/ (2024) (2007).

23. Partridge, A. W., Therien, A. G. & Deber, C. M. Polar mutations in membrane proteins as a biophysical basis for disease. Peptide Science: Original Research on Biomolecules 66, 350–358 (2002).

24. Flagmeier, P. et al. Mutations associated with familial Parkinson’s disease alter the initiation and amplification steps of α-synuclein aggregation. Proceedings of the National Academy of Sciences 113, 10328–10333 (2016).

25. Nagy, J. K. & Sanders, C. R. Destabilizing mutations promote membrane protein misfolding. Biochemistry 43, 19–25 (2004).

26. Lei, X. et al. The cancer mutation D83V induces an α-helix to β-strand conformation switch in MEF2B. Journal of molecular biology 430, 1157–1172 (2018).

27. Anderson, T. A., Cordes, M. H. J. & Sauer, R. T. Sequence determinants of a conformational switch in a protein structure. Proceedings of the National Academy of Sciences 102, 18344–18349. eprint: 10.1073/pnas.0509349102. https://www.pnas.org/doi/abs/10.1073/pnas.0509349102(2005).

28. Chakravarty, D., Sreenivasan, S., Swint-Kruse, L. & Porter, L. L. Identification of a covert evolutionary pathway between two protein folds. Nature Communications 14, 3177. issn: 2041-1723. 10.1038/s41467-023-38519-0(June 2023).

29. Abrusn, G. & Marsh, J. A. Alpha Helices Are More Robust to Mutations than Beta Strands. PLoS Computational Biology 12, e1005242. issn: 1553-734X. https://www.ncbi.nlm.nih.gov/pmc/articles/PMC5147804/ (2023) (Dec. 2016).

30. Greenfield, N. J. Using circular dichroism spectra to estimate protein secondary structure. Nature protocols 1, 2876–2890. issn: 1754-2189. https://www.ncbi.nlm.nih.gov/pmc/articles/PMC2728378/ (2024) (2006).

31. Hermans, J. Molecular dynamics simulations of helix and turn propensities in model peptides. Current Opinion in Structural Biology 3, 270–276 (1993).

32. Wu, R. et al. High-resolution de novo structure prediction from primary sequence. bioRxiv. eprint: https://www.biorxiv.org/content/early/2022/07/22/2022.07.21.500999.full.pdf. https://www.biorxiv.org/content/early/2022/07/22/2022.07.21.500999 (2022).

33. Kabsch, W. & Sander, C. Dictionary of protein secondary structure: pattern recognition of hydrogen-bonded and geometrical features. Biopolymers: Original Research on Biomolecules 22, 2577–2637 (1983).

34. Labesse, G., Colloc’h, N., Pothier, J. & Mornon, J.-P. P-SEA: a new efficient assignment of secondary structure from Cα trace of proteins. Bioinformatics 13, 291–295 (1997).

35. Lyu, Z., Wang, Z., Luo, F., Shuai, J. & Huang, Y. Protein secondary structure prediction with a reductive deep learning method. Frontiers in Bioengineering and Biotechnology 9, 687426 (2021).

36. Yu, C.-H. et al. End-to-end deep learning model to predict and design secondary structure content of structural proteins. ACS biomaterials science & engineering 8, 1156–1165 (2022).

37. Sutton, R. & Barto, A. Reinforcement Learning: An Introduction isbn: 9780262193986. https://books.google.co.in/books?id=CAFR6IBF4xYC (MITPress. 1998).

38. Wang, Y. et al. Self-play reinforcement learning guides protein engineering. Nature Machine Intelligence 5, 845–860 (2023).

39. Angermueller, C. et al. Model-based reinforcement learning for biological sequence design in International conference on learning representations (2019).

40. Silver, D. et al. Mastering the game of Go without human knowledge. en. Nature 550, 354–359. issn: 1476-4687. https://www.nature.com/articles/nature24270(2024) (Oct. 2017).

41. Silver, D. et al. A general reinforcement learning algorithm that masters chess, shogi, and Go through self-play. en. Science 362, 1140–1144. issn: 0036-8075, 1095-9203. 10.1126/science.aar6404 (2024) (Dec.2018).

42. Williams, R. J. Simple statistical gradient-following algorithms for connectionist reinforcement learning. Machine learning 8, 229–256 (1992).

43. Mnih, V. et al. Asynchronous Methods for Deep Reinforcement Learning 2016.1602.01783 [cs.LG]. https://arxiv.org/abs/1602.01783.

44. Haarnoja, T., Zhou, A., Abbeel, P. & Levine, S. Soft Actor-Critic: Off-Policy Maximum Entropy Deep Reinforcement Learning with a Stochastic Actor 2018. 1801.01290 [cs.LG]. https://arxiv.org/abs/1801.01290.

45. Schulman, J., Wolski, F., Dhariwal, P., Radford, A. & Klimov, O. Proximal Policy Optimization Algorithms 2017. 1707.06347 [cs.LG]. https://arxiv.org/abs/1707.06347.

46. Mnih, V. et al. Human-level control through deep reinforcement learning. en. Nature 518, 529–533. issn: 1476-4687. https://www.nature.com/articles/nature14236 (2024) (Feb. 2015).

47. Asgari, E. & Mofrad, M. R. Continuous distributed representation of biological sequences for deep proteomics and genomics. PloS one 10, e0141287 (2015).

48. Towers, M. et al. Gymnasium Mar. 2023. https://zenodo.org/record/8127025 (2023).

49. Tsai, C.-Y. et al. Helical structure motifs made searchable for functional peptide design. en. Nature Communications 13. Number: 1 Publisher: Nature Publishing Group, 102. issn: 2041-1723. https://www.nature.com/articles/s41467-021-27655-0 (2023) (Jan. 2022).

50. Goodsell, D. S. et al. RCSB Protein Data Bank: Enabling biomedical research and drug discovery. en. Protein Science 29, 52–65. issn: 1469-896X. 10.1002/pro.3730 (2023) (2020).

51. Raffin, A. et al. Stable-Baselines3: Reliable Reinforcement Learning Implementations. Journal of Machine Learning Research 22, 1–8. http://jmlr.org/papers/v22/20-1364.html (2021).

52. Eastman, P. et al. OpenMM 7: Rapid development of high performance algorithms for molecular dynamics. PLoS computational biology 13, e1005659 (2017).

53. Izadi, S. & Onufriev, A. V. Accuracy limit of rigid 3-point water models. The Journal of chemical physics 145 (2016).

54. Maier, J. A. et al. ff14SB: improving the accuracy of protein side chain and backbone parameters from ff99SB. Journal of chemical theory and computation 11, 3696–3713 (2015).

55. Bussi, G., Donadio, D. & Parrinello, M. Canonical sampling through velocity rescaling. The Journal of chemical physics 126 (2007).

56. Parrinello, M. & Rahman, A. Polymorphic transitions in single crystals: A new molecular dynamics method. Journal of Applied physics 52, 7182–7190 (1981).

57. Abraham, M. J. et al. GROMACS: High performance molecular simulations through multi-level parallelism from laptops to supercomputers. SoftwareX 1, 19–25 (2015).

58. Tribello, G. A., Bonomi, M., Branduardi, D., Camilloni, C. & Bussi, G. PLUMED 2: New feathers for an old bird. Computer physics communications 185, 604–613 (2014).

59. Singh, R. K., Chamachi, N. G., Chakrabarty, S. & Mukherjee, A. Mechanism of Unfolding of Human Prion Protein. The Journal of Physical Chemistry B 121, 550–564. issn: 1520-6106. 10.1021/acs.jpcb.6b11416 (2024) (Jan. 26, 2017).

60. Negrete, J. A., Viñuales, Y. & Palau, J. Deciphering the structural code for proteins: Helical propensities in domain classes and statistical multiresidue information in -helices. en. Protein Science 7, 1368–1379. issn: 0961-8368, 1469-896X. 10.1002/pro.5560070613 (2024) (June 1998).

61. Bhattacharjee, N. & Biswas, P. Helical ambivalency induced by point mutations. BMC Structural Biology 13, 1–11 (2013).

62. Myers, J. K., Pace, C. N. & Scholtz, J. M. Helix Propensities Are Identical in Proteins and Peptides. Biochemistry 36. Publisher: American Chemical Society, 10923–10929. issn: 0006-2960. 10.1021/bi9707180 (2024) (Sept. 1997).

63. Gellera, C. et al. Frataxin gene point mutations in Italian Friedreich ataxia patients. en. Neurogenetics 8, 289–299. issn: 1364-6753. 10.1007/s10048-007-0101-5 (2024) (Nov. 2007).

64. Clark, E., Butler, J. S., Isaacs, C. J., Napierala, M. & Lynch, D. R. Selected missense mutations impair frataxin processing in Friedreich ataxia. Annals of Clinical and Translational Neurology 4, 575–584. issn: 2328-9503. https://www.ncbi.nlm.nih.gov/pmc/articles/PMC5553228/ (2024) (June 2017).

65. Zheng, Z., Chinnasamy, N. & Morgan, R. A. Protein L: a novel reagent for the detection of Chimeric Antigen Receptor (CAR) expression by flow cytometry. Journal of Translational Medicine 10, 29. issn: 1479-5876. 10.1186/1479-5876-10-29 (2024) (Feb. 2012).

66. Da Conceição, L. M. A., Cabral, L. M., Pereira, G. R. C. & De Mesquita, J. F. An In Silico Analysis of Genetic Variants and Structural Modeling of the Human Frataxin Protein in Friedreich’s Ataxia. en. International Journal of Molecular Sciences 25. Number: 11 Publisher: Multidisciplinary Digital Publishing Institute, 5796. issn: 1422-0067. https://www.mdpi.com/1422-0067/25/11/5796 (2024) (Jan. 2024).

67. Botticelli, S. et al. Modelling Protein Plasticity: The Example of Frataxin and Its Variants. Molecules 27, 1955. issn: 1420-3049. https://www.ncbi.nlm.nih.gov/pmc/articles/PMC8950120/ (2024) (Mar. 2022).

68. Kim, D. E., Fisher, C. & Baker, D. A breakdown of symmetry in the folding transition state of protein L1. Journal of Molecular Biology 298, 971– 984. issn: 0022-2836. https://www.sciencedirect.com/science/article/pii/S002228360093701X (2024) (May 2000).

69. Webb, B. & Sali, A. Comparative Protein Structure Modeling Using _e_print. en. Current Protocols in Bioinformatics 54, 5.6.1–5.6.37. issn: 1934-340X. 10.1002/cpbi.3 (2023) (2016).

70. Cheng, J. et al. Accurate proteome-wide missense variant effect prediction with AlphaMissense. Science 381, eadg7492 (2023).

71. Ittisoponpisan, S. et al. Can predicted protein 3D structures provide reliable insights into whether missense variants are disease associated? Journal of molecular biology 431, 2197–2212 (2019).

72. Dieckhaus, H., Brocidiacono, M., Randolph, N. Z. & Kuhlman, B. Transfer Learning to Leverage Larger Datasets for Improved Prediction of Protein Stability Changes. Proceedings of the National Academy of Sciences 121, e2314853121. 10.1073/pnas.2314853121 (2024) (Feb. 6, 2024).

73. Blaabjerg, L. M. et al. Rapid Protein Stability Prediction Using Deep Learning Representations. eLife 12 (eds Faraldo-Gómez, J. D., Weigel, D., Ben-Tal, N. & Echave, J.) e82593. issn: 2050-084X. 10.7554/eLife.82593 (2024) (May 15, 2023).

74. Cao, H., Wang, J., He, L., Qi, Y. & Zhang, J. Z. DeepDDG: Predicting the Stability Change of Protein Point Mutations Using Neural Networks. Journal of Chemical Information and Modeling 59, 1508–1514. issn: 1549-9596. 10.1021/acs.jcim.8b00697 (2024) (Apr. 22, 2019).

