## Supporting Information for "AlphaMut: a deep reinforcement learning model to suggest helix-disrupting mutations"

### Choosing the appropriate reinforcement learning algorithm

A2C, DQN and PPO algorithms were implemented with default parameters from the StableBaselines3 implementation, for 168000 episodes. Convergence plots are shown in Figure S1.

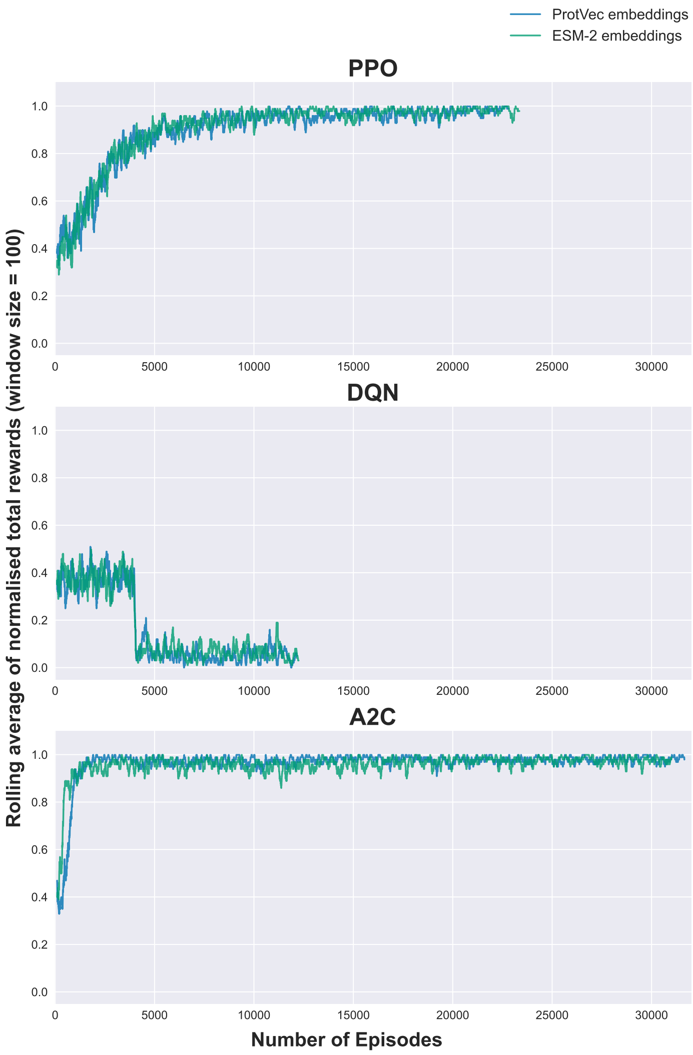


Figure S1: Convergence plots for different algorithms for the Helix-only model with different algorithms, Actor to Critic (A2C), Deep Q Networks (DQN) and Proximal Policy Optimisation (PPO).

### Validation metrics for the Helix-only model

$\beta$ and $\gamma$ were varied for the different algorithms. These correspond to the `ent_coef` and `gamma` respectively in the package. The validation metrics calculated for the trained models of A2C and DQN are provided in Table S1 and for PPO in Table S2.

Table S1: Validation metrics for DQN and PPO runs for the Helix-only model.

| **A2C** | | | | | |
|---|---|---|---|---|---|
| proline | $\beta$ | $\gamma$ | Learning Efficiency | Mutational Entropy (S) | Embeddings |
| True | 0 | 0.99 | 0.98 | 0.01 | ProtVec |
| True | 0 | 0.99 | 0.97 | 0.28 | ESM-2 |
| False | 0 | 0.99 | 0.2 | 2.2 | ProtVec |
| False | 0 | 0.99 | 0.23 | 2.4 | ESM-2 |
| **DQN** | | | | | |
| proline | $\beta$ | $\gamma$ | Learning Efficiency | Mutational Entropy (S) | Embeddings |
| True | - | 0.99 | 0.06 | 1.35 | ProtVec |
| True | - | 0.99 | 0.03 | 1.41 | ESM-2 |
| False | - | 0.99 | 0.04 | 1.85 | ProtVec |
| False | - | 0.99 | 0.05 | 1.51 | ESM-2 |

Table S2: Validation metrics for the Helix-only models trained using the PPO algorithm.

| **PPO** | | | | |
|---|---|---|---|---|
| **With proline** | | | | |
| Embedding | $\beta$ | $\gamma$ | Learning Efficiency ($\kappa$) | Mutation Entropy ($S$) |
| ProtVec | 0.001 | 0.95 | 0.99 | 0.85 |
| ESM-2 | 0.001 | 0.95 | 0.99 | 0.69 |
| ProtVec | 0 | 0.99 | 0.98 | 1.89 |
| ESM-2 | 0 | 0.99 | 0.98 | 1.63 |
| **Without proline** | | | | |
| Embedding | $\beta$ | $\gamma$ | Learning Efficiency ($\kappa$) | Mutation Entropy ($S$) |
| ProtVec | 0.001 | 0.95 | 0.81 | 2.19 |
| ESM-2 | 0.001 | 0.95 | 0.84 | 2.17 |
| ProtVec | 0 | 0.99 | 0.84 | 2.14 |
| ESM-2 | 0 | 0.99 | 0.86 | 2.03 |

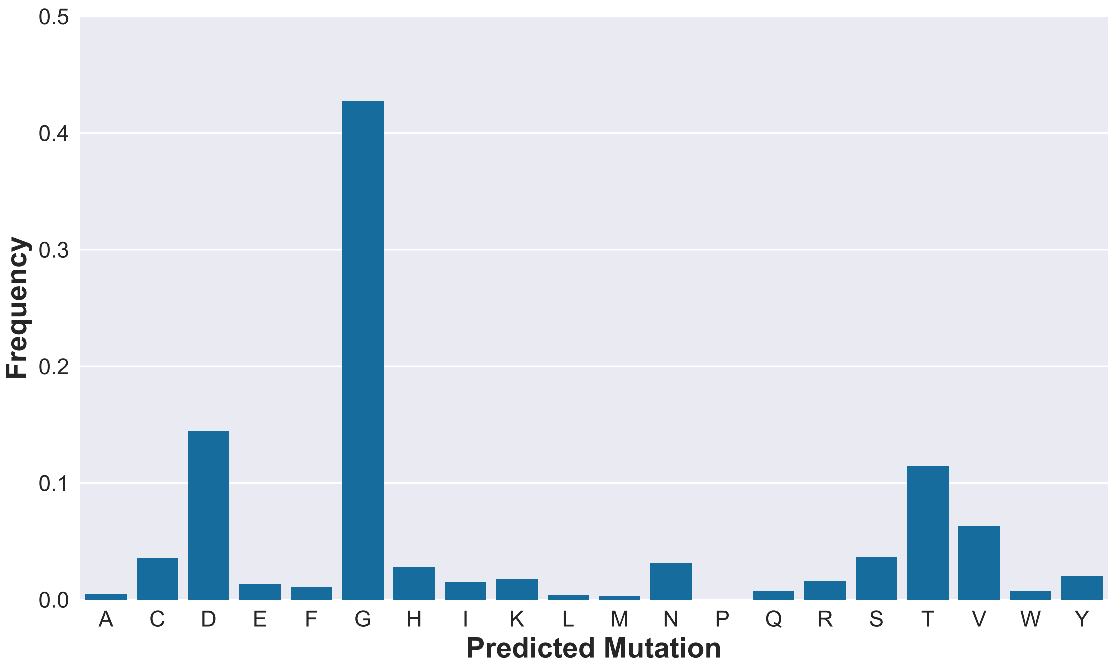


Figure S2: Frequencies of predicted mutations when the Helix-only model is trained on the TP dataset without proline as an allowed mutation. The model chosen is the embedded with ESM-2 with hyperparameters $\beta = 0.001$ and $\gamma = 0.95$, highlighted in green in Table S2.

| Embedding | β | γ | Learning Efficiency | Mutation Entropy |
|---|---|---|---|---|
| ProtVec | 0.001 | 0.95 | 0.99 | 0.85 |
| ESM-2 | 0.001 | 0.95 | 0.99 | 0.69 |
| ProtVec | 0 | 0.99 | 0.98 | 1.89 |
| ESM-2 | 0 | 0.99 | 0.98 | 1.63 |

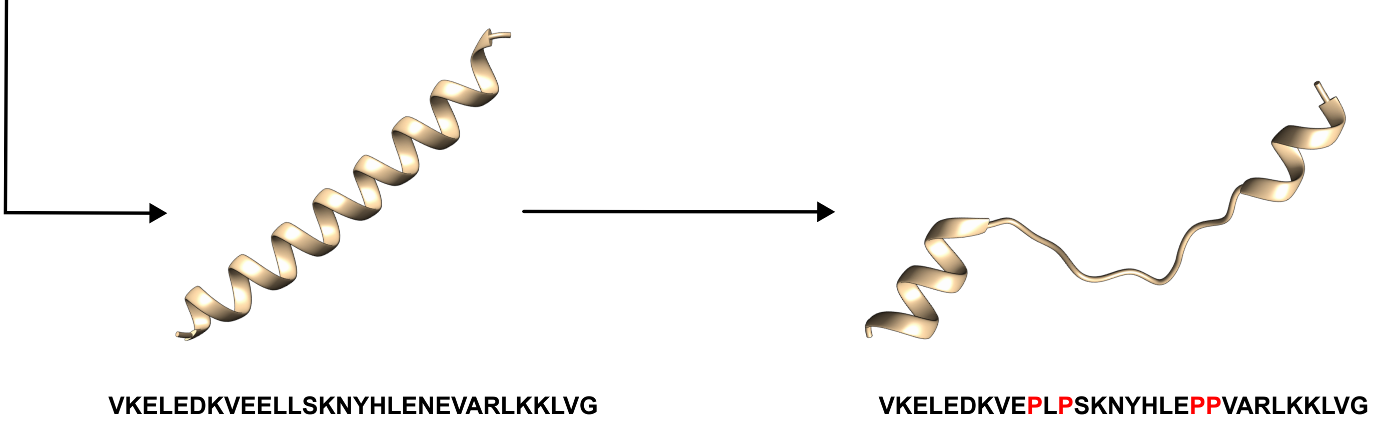


Figure S3: Validation metrics of different evaluated models, with the chosen model for disruption highlighted in orange. Also shown is a disrupted structure obtained from running the model on helix capping in the GCN4 leucine zipper protein (PDB ID: 1ce9), with residues 3-32.

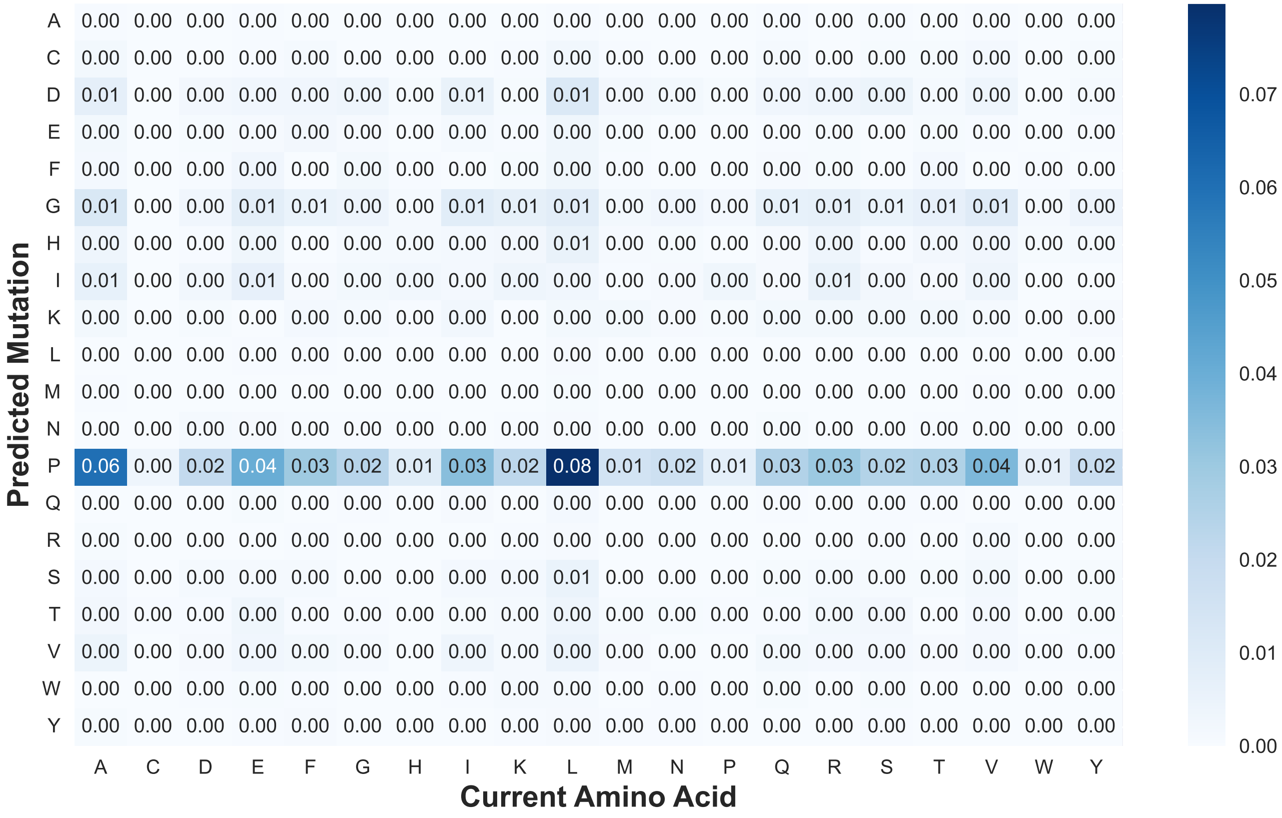


Figure S4: Heatmap of current amino acids against predicted mutations by the chosen model, highlighted in pink in Table S2.

### Convergence plots for the Helix-in-protein model

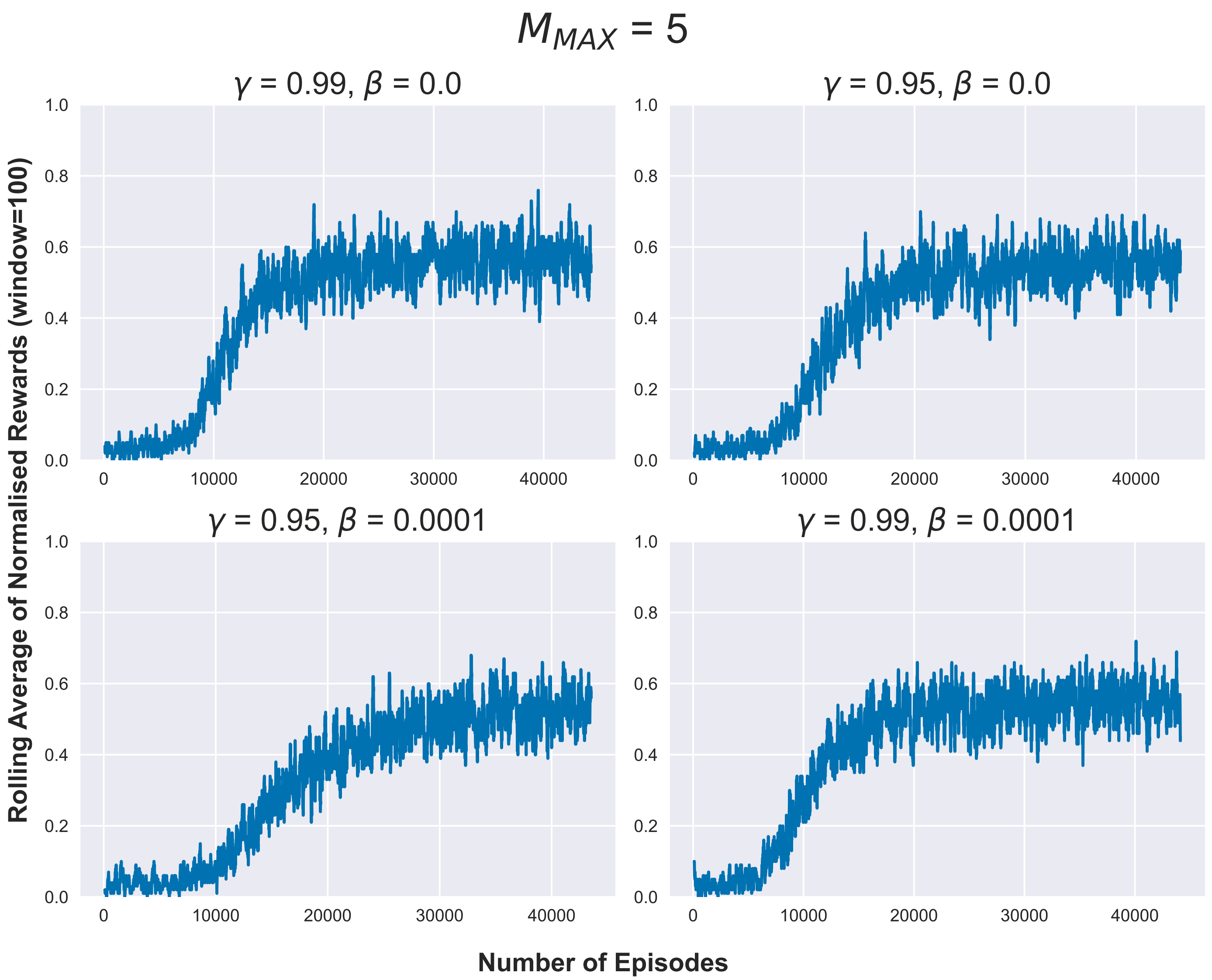


Figure S5: Convergence plots for PPO training with $M_{MAX} = 5$ for the Helix-in-protein model.

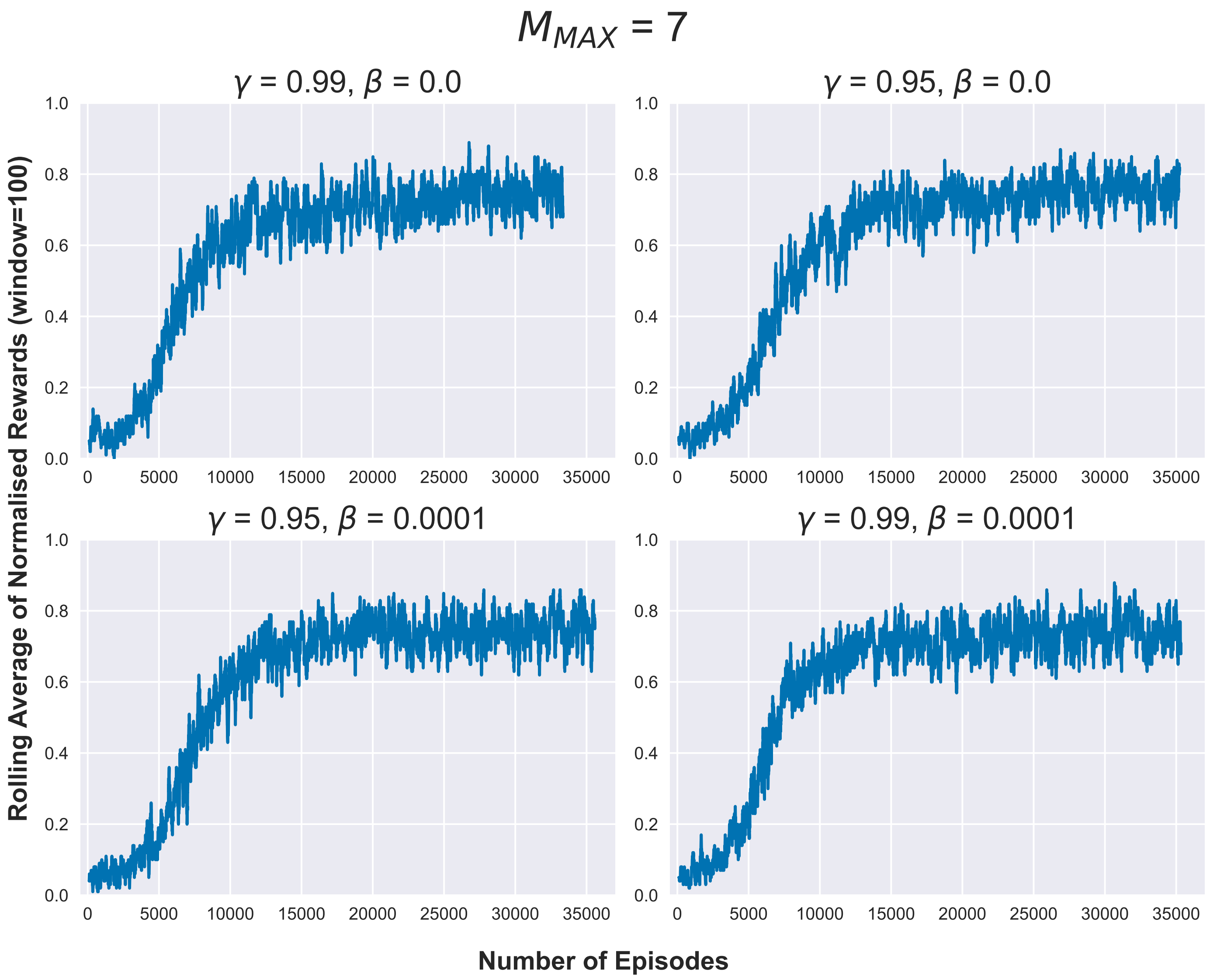


Figure S6: Convergence plots for PPO training with $M_{MAX} = 7$ for the Helix-in-protein model.

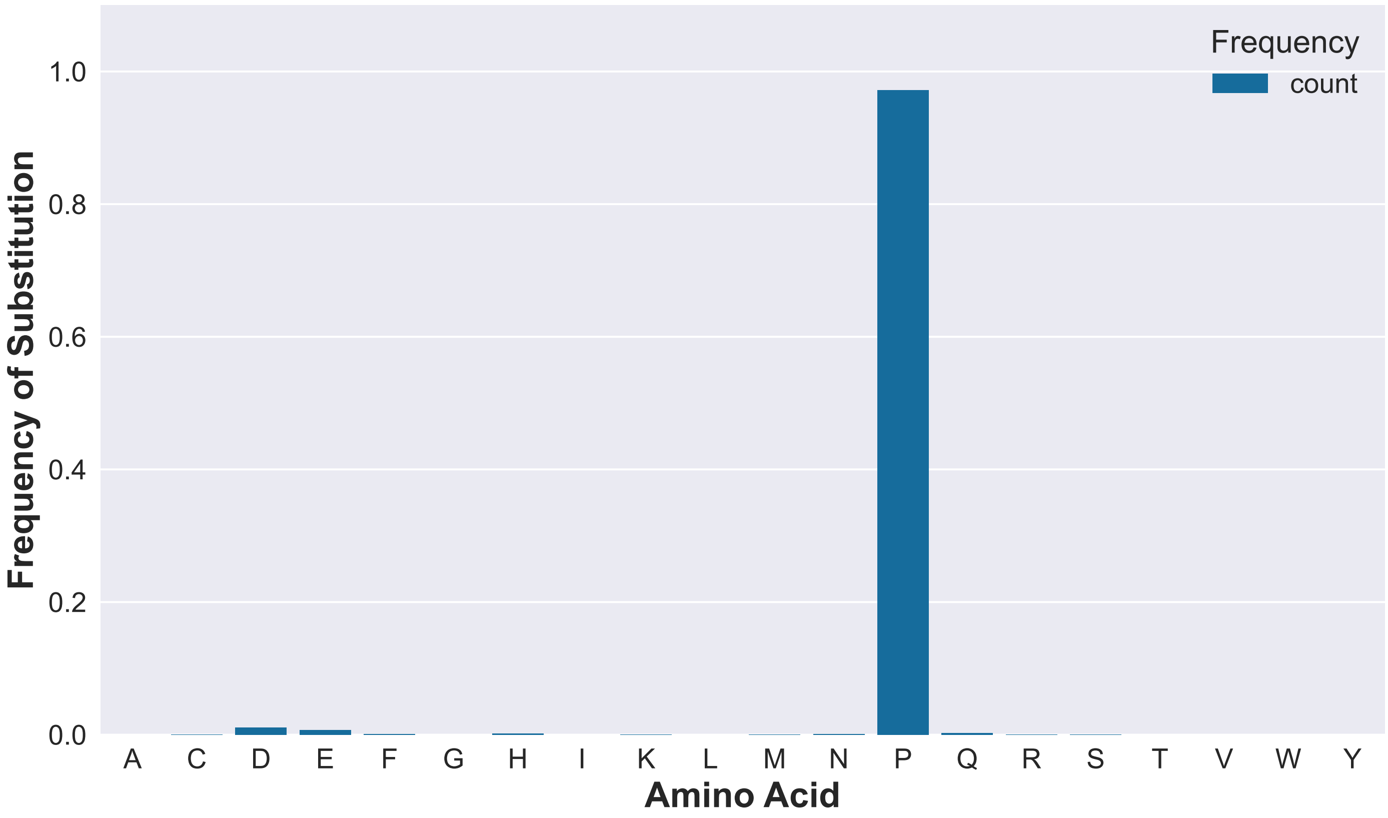


Figure S7: Substitution frequencies for $\gamma = 0.95$ and $\beta = 0.0001$ with $M_{MAX} = 5$.

### Validation metrics for Helix-in-protein model

Table S3: Validation metrics for Helix-in-protein model.

| $M_{MAX} = 5$ | | | | |
|---|---|---|---|---|
| $\beta$ | $\gamma$ | Learning Efficiency ($\kappa$) | Mutation Entropy ($S$) | Average Number of Mutations |
| 0.0 | 0.95 | 0.56 | 0.08 | 3.59 |
| 0.0 | 0.99 | 0.54 | 0.1 | 3.62 |
| 0.0001 | 0.99 | 0.56 | 0.08 | 3.65 |
| 0.0001 | 0.95 | 0.53 | 0.18 | 3.48 |
| $M_{MAX} = 7$ | | | | |
| $\beta$ | $\gamma$ | Learning Efficiency ($\kappa$) | Mutation Entropy ($S$) | Average Number of Mutations |
| 0.0 | 0.95 | 0.75 | 0.31 | 4.46 |
| 0.0 | 0.95 | 0.75 | 0.31 | 4.46 |
| 0.0001 | 0.99 | 0.77 | 0.34 | 4.5 |
| 0.0001 | 0.95 | 0.74 | 0.39 | 4.53 |

### Comparison of propensities between Helix-only model and Helix-in-protein model

Propensities calculated from the Helix-only and the Helix-in-protein model were compared and a Spearmann coefficient $\rho = 0.43$ was obtained. Propensity obtained from our Helix-in-protein model correlates best with Moreau and coworkers[1] with a $\rho$ of 0.45. Myers and coworkers[2] ranks matches best with our index from the Helix-only model, with a $\rho = 0.54$. Notably, both these propensity indices are from experimental studies shaded in blue in Table S4. The propensity indices obtained from bioinformatics studies are shaded in brown.

Table S4: Comparison of propensity scales from different studies (cited in the corresponding column headers) and their Spearmann correlation coefficients ($\rho$) with the propensity scales from our Helix-only and Helix-in-protein models. The cells corresponding to the best $\rho$ for each propensity scale are highlighted in purple.

| Helix-only | Helix-in-protein | [3] | [1] | [4] | [5] | [6] | [7] | [5] | [2] | [8] | [2] | [9] | [10] |
|---|---|---|---|---|---|---|---|---|---|---|---|---|---|
| L | I | A | A | A | A | A | A | A | N | A | A | R | A |
| A | L | W | L | L | L | E | E | L | A | R | W | A | R |
| R | R | E | E | M | R | M | L | M | L | K | M | K | K |
| Q | K | L | M | I | M | L | M | R | M | M | L | L | L |
| M | A | Q | I | Q | K | Q | Q | K | W | L | Q | E | M |
| E | E | Y | R | R | Q | R | R | Q | Q | S | E | M | Q |
| Y | M | K | Q | K | E | W | K | E | Y | Q | I | F | E |
| N | T | M | W | Y | I | K | W | I | R | E | Y | W | Y |
| F | V | R | K | V | W | F | I | S | I | N | K | Q | I |
| S | S | F | V | F | S | Y | F | W | S | F | S | I | S |
| K | F | I | D | W | Y | D | Y | Y | K | D | R | Y | D |
| V | N | H | F | H | F | C | V | F | H | H | C | C | C |
| T | Y | D | Y | T | V | I | H | V | F | T | F | S | W |
| W | Q | N | S | E | H | H | C | T | T | I | N | D | N |
| I | W | T | N | S | N | V | D | H | V | Y | V | N | F |
| D | G | V | T | D | T | T | T | C | E | V | T | H | V |
| C | H | C | C | C | C | N | S | N | D | G | D | V | H |
| H | C | S | G | N | D | S | N | D | C | W | G | T | T |
| G | D | G | H | G | G | P | P | G | G | C | P | G | G |
| P | P | P | P | P | P | G | G | P | P | P | H | P | P |
| $\rho$ (Helix-only) | 0.43 | -0.43 | 0.28 | -0.02 | -0.06 | -0.03 | -0.21 | 0.17 | **0.54** | -0.17 | -0.16 | -0.22 | -0.1 |
| $\rho$ (Helix-in-protein) | 1.00 | -0.17 | **0.45** | 0.27 | 0.34 | -0.12 | -0.44 | 0.33 | 0.22 | 0.11 | 0.15 | 0.09 | 0.13 |

### Convergence plot for the Helix-in-protein model without proline

The Helix-in-protein model without proline as an allowed mutation was trained for 168000 steps with $\beta = 0.0$ and $\gamma = 0.99$. The convergence plot for the same is shown in Figure S8.

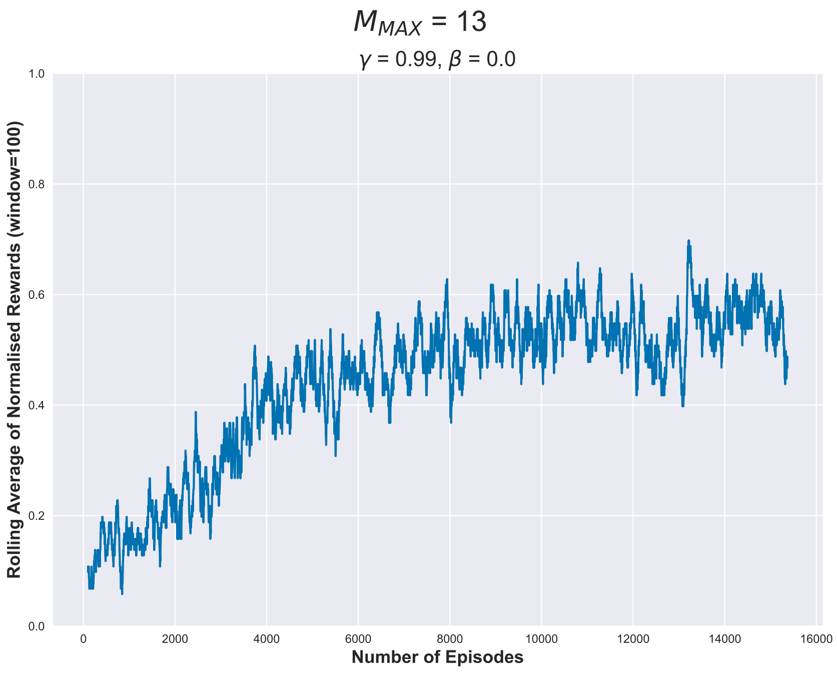


Figure S8: Convergence plot for training of Helix-in-protein environment without proline.

### ESMFold predicted structures of wild-type and mutant proteins for mutations predicted by our model

In this section, we provide the ESMFold predicted mutated structures of some proteins of interest. The green shade indicates the residues that are part of the helix in the wild-type structure. The wild-type reference structures are provided in the right corner of each figure, with the PDB IDs for each protein provided in Table S5. The starting and ending residue IDs for the helices chosen for disruption are also provided in the same table.

Table S5: Wild-type PDB IDs for chosen proteins.

| **protein** | **PDB ID** | **chain** | **starting residue ID** | **ending residue ID** |
|---|---|---|---|---|
| insulin | 1lph | D | 4 | 18 |
| zinc finger protein | 1wjp | A | 28 | 42 |
| haemoglobin | 5hbi | A | 29 | 43 |
| myoglobin | 1do4 | A | 82 | 96 |
| HSP-90 | 2yi0 | A | 110 | 124 |

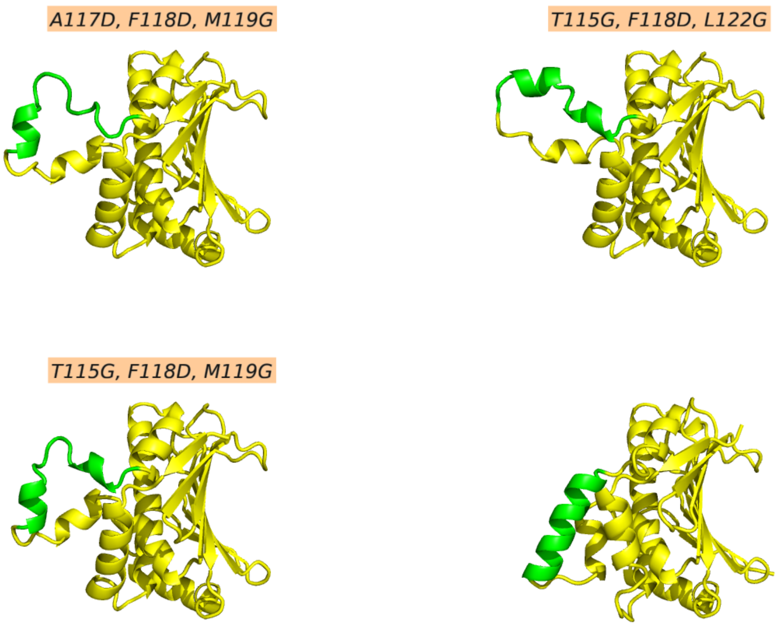


Figure S9: Wild-type(right corner) and mutant structures with indicated mutations in orange for HSP-90.

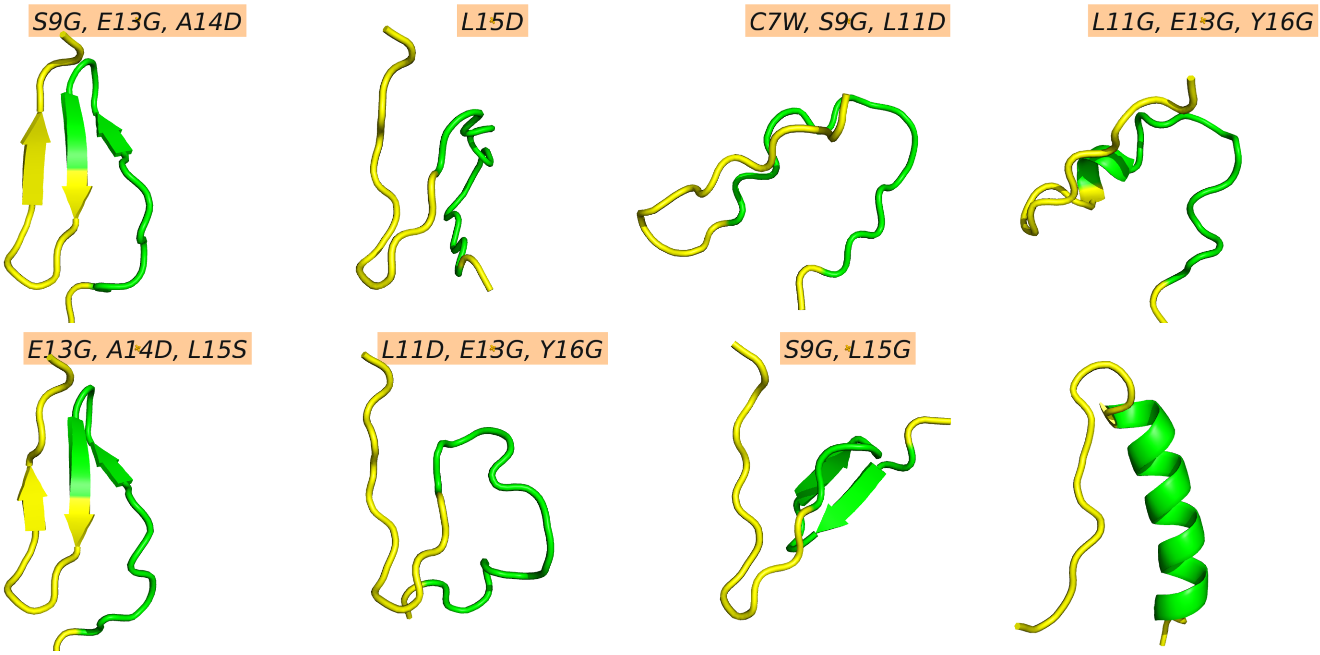


Figure S10: Wild-type(right corner) and mutant structures with indicated mutations in orange for insulin.

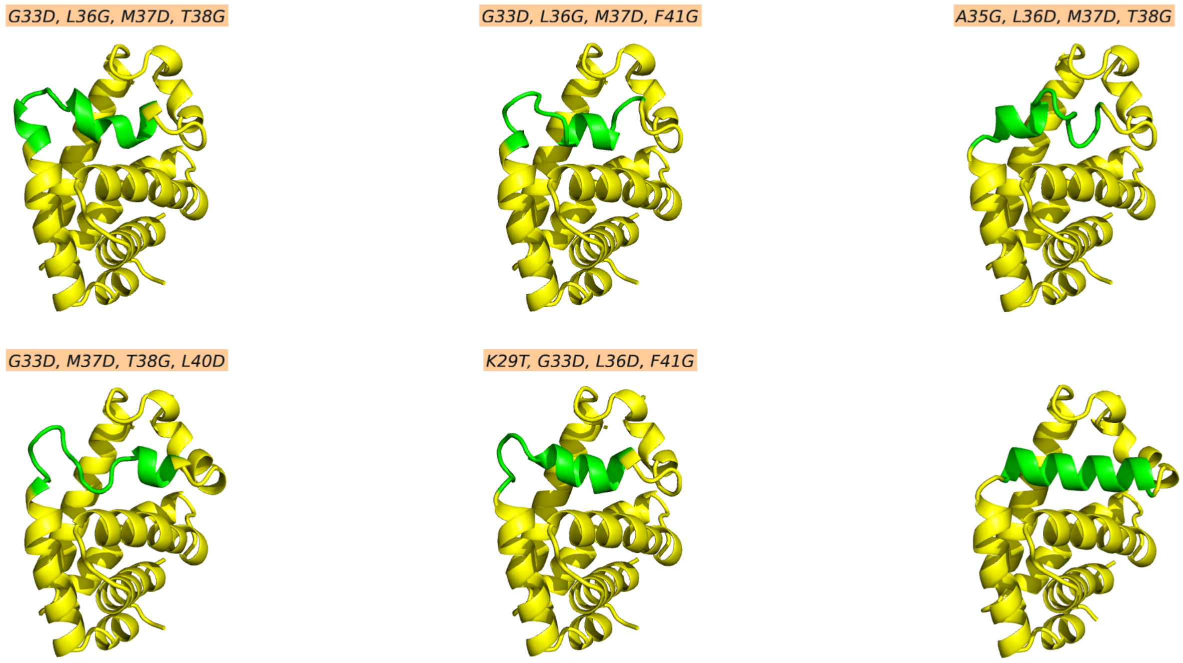


Figure S11: Wild-type(right corner) and mutant structures with indicated mutations in orange for haemoglobin.

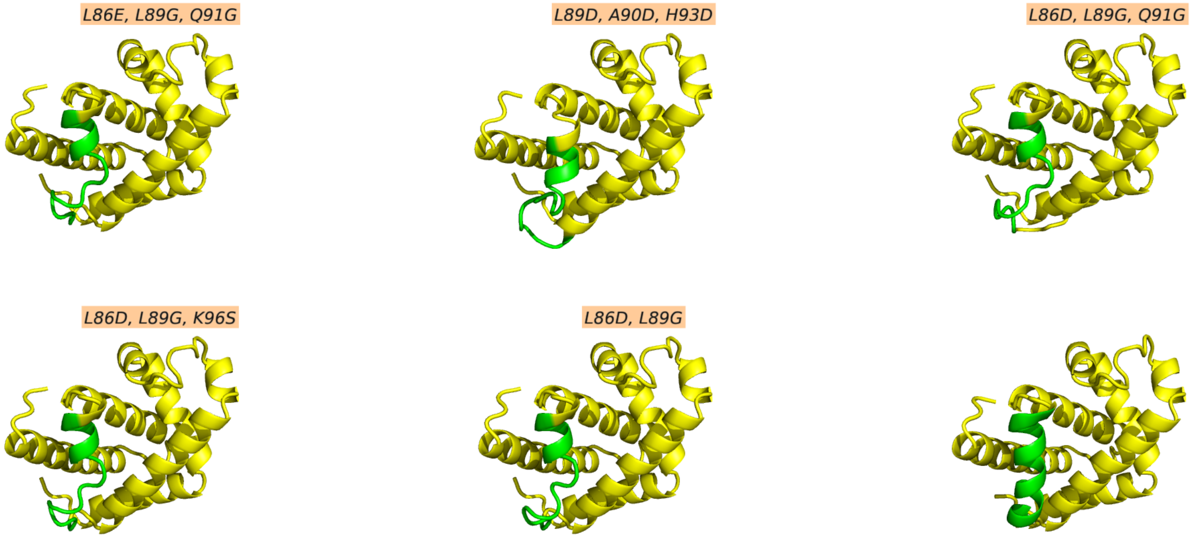


Figure S12: Wild-type(right corner) and mutant structures with indicated mutations in orange for myoglobin.

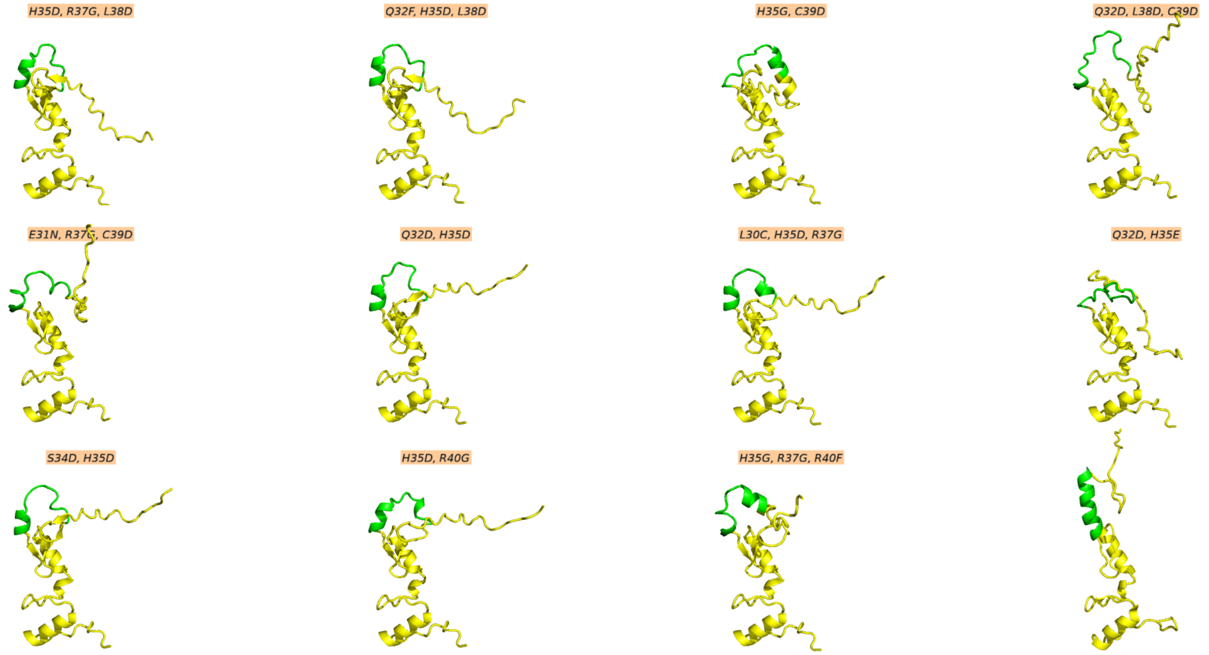


Figure S13: Wild-type(right corner) and mutant structures with indicated mutations in orange for zinc finger protein.

### Metadynamics simulation details for validation of Helix-in-protein model

To determine the appropriate hill width value, we conducted brief 10 ns unbiased simulations. The values of hill width were selected based on one-half of the standard deviation of the collective variable $\langle\Psi\rangle$ obtained from the unbiased simulations. We executed well-tempered metadynamics simulations with the parameters given below for the four systems in Table S6.

Table S6: Parameters for well-tempered metadynamics simulations of frataxin and protein-L systems.

| **System** | **Hill Width** $(Radian)$ | **Hill Height** $(kJ/mol)$ | **BIASFACTOR** | **PACE** $(ps)$ |
|---|---|---|---|---|
| frataxin wild-type | 0.02 | 0.8 | 8 | 500 |
| frataxin mutant | 0.03 | 0.5 | 8 | 1000 |
| protein-L wild-type | 0.02 | 0.5 | 8 | 1000 |
| protein-L mutant | 0.02 | 0.5 | 8 | 1000 |

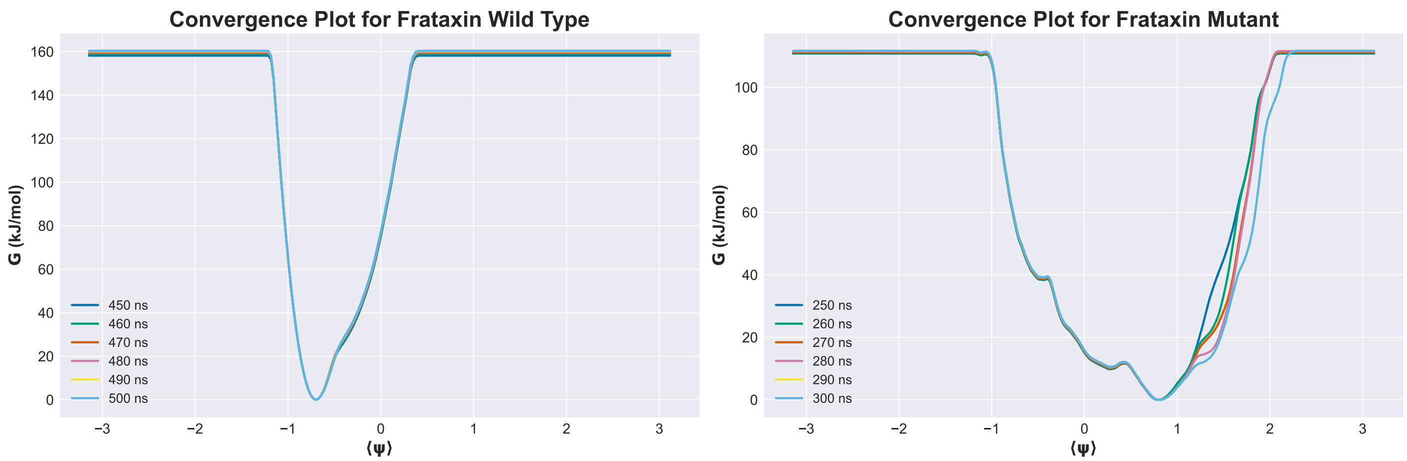


Figure S14: Convergence plots for the wild-type and mutant well-tempered metadynamics simulations of frataxin.

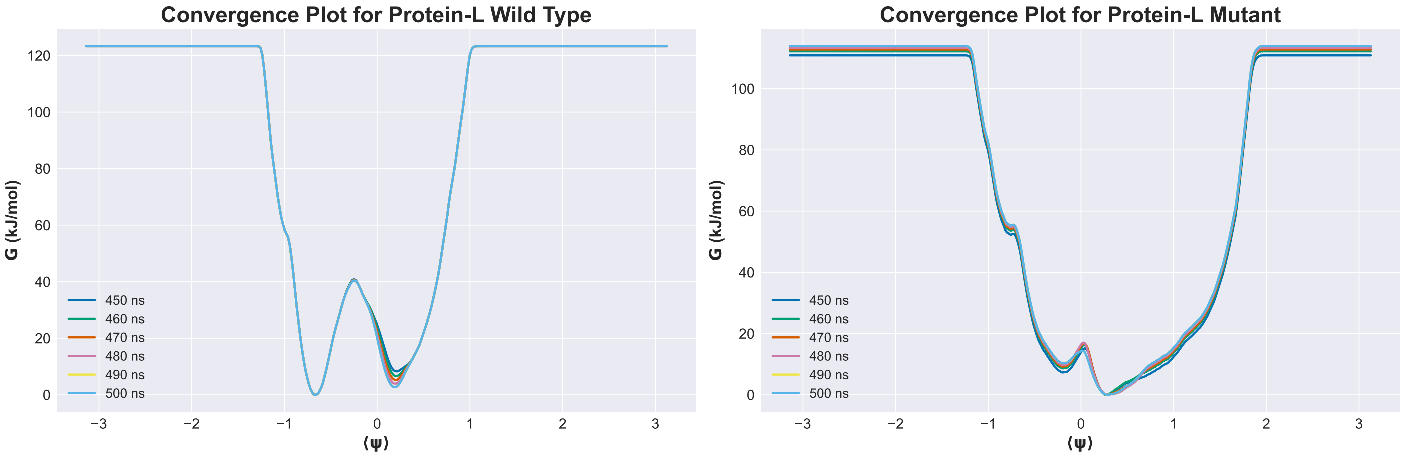


Figure S15: Convergence plots for the wild-type and mutant well-tempered metadynamics simulations of protein-L.

### Bibliography


1. Moreau, R. J. *et al.* Context-Independent, Temperature-Dependent Helical Propensities for Amino Acid Residues. *Journal of the American Chemical Society* **131.** PMID: 19702302, 13107–13116. eprint: https://doi.org/10.1021/ja904271k. https://doi.org/10.1021/ja904271k (2009).
2. Myers, J. K., Pace, C. N. & Scholtz, J. M. Helix Propensities Are Identical in Proteins and Peptides. *Biochemistry* **36.** Publisher: American Chemical Society, 10923–10929. ISSN: 0006-2960. https://doi.org/10.1021/bi9707180 (2024) (Sept. 1997).
3. Tsai, C.-Y. *et al.* Helical structure motifs made searchable for functional peptide design. en. *Nature Communications* **13.** Number: 1 Publisher: Nature Publishing Group, 102. ISSN: 2041-1723. https://www.nature.com/articles/s41467-021-27655-0 (2023) (Jan. 2022).
4. Blaber, M., Zhang, X.-j. & Matthews, B. W. Structural basis of amino acid $\alpha$ helix propensity. *Science* **260,** 1637–1640 (1993).
5. Pace, C. N. & Scholtz, J. M. A helix propensity scale based on experimental studies of peptides and proteins. *Biophysical journal* **75,** 422–427 (1998).
6. Shingate, P. & Sowdhamini, R. Analysis of domain-swapped oligomers reveals local sequence preferences and structural imprints at the linker regions and swapped interfaces. *PloS one* (2012).
7. Fujiwara, K., Toda, H. & Ikeguchi, M. Dependence of $\alpha$-helical and $\beta$-sheet amino acid propensities on the overall protein fold type. *BMC structural biology* **12,** 1–15 (2012).
8. Horovitz, A., Matthews, J. M. & Fersht, A. R. -Helix stability in proteins: II. Factors that influence stability at an internal position. *Journal of Molecular Biology* **227,** 560–568. ISSN: 0022-2836. https://www.sciencedirect.com/science/article/pii/0022283692909072 (1992).
9. Park, S. H., Shalongo, W. & Stellwagen, E. Residue helix parameters obtained from dichroic analysis of peptides of defined sequence. *Biochemistry* **32.** PMID: 8334134, 7048–7053. eprint: https://doi.org/10.1021/bi00078a033. https://doi.org/10.1021/bi00078a033 (1993).
10. Rohl, C. A., Chakrabartty, A. & Baldwin, R. L. Helix propagation and N-cap propensities of the amino acids measured in alanine-based peptides in 40 volume percent trifluoroethanol. *Protein Science* **5,** 2623–2637 (1996).
